# Transcriptomic signatures of progressive and regressive liver fibrosis and portal hypertension

**DOI:** 10.1101/2023.07.22.550133

**Authors:** Oleksandr Petrenko, Philipp Königshofer, Ksenia Brusilovskaya, Benedikt S Hofer, Katharina Bareiner, Benedikt Simbrunner, Michael Trauner, Stefan G Kauschke, Larissa Pfisterer, Eric Simon, André F Rendeiro, Laura P M H de Rooij, Philipp Schwabl, Thomas Reiberger

**Affiliations:** Division of Gastroenterology and Hepatology, Department of Internal Medicine III, Medical University of Vienna, Vienna, Austria; Christian Doppler Laboratory for Portal Hypertension and Liver Fibrosis, Medical University of Vienna, Vienna, Austria; CeMM Research Center for Molecular Medicine of the Austrian Academy of Sciences, Vienna, Austria; Department of CardioMetabolic Diseases Research, Boehringer Ingelheim Pharma GmbH & Co.KG, Biberach an der Riss, Germany; Global Computational Biology and Digital Sciences, Boehringer Ingelheim Pharma GmbH & Co.KG, Biberach an der Riss, Germany

**Author notes:** Correspondence should be addressed to T.R.

## Abstract

Persistent liver injury triggers a fibrogenic program that causes pathologic remodelling of the hepatic microenvironment (i.e., liver fibrosis) and portal hypertension. The dynamics of gene regulation during liver disease progression and regression remain understudied. Here, we generated hepatic transcriptome profiles in two well-established liver disease models at peak fibrosis and during spontaneous regression after the removal of the inducing agents. We linked the dynamics of key liver disease readouts, such as portal pressure, collagen proportionate area, and transaminase serum levels, to most differentially expressed genes, enabling the identification of transcriptomic signatures of progressive vs. regressive liver fibrosis and portal hypertension. These candidate biomarkers (e.g., *Scube1*, *Tcf4*, *Src*, *Hmga1*, *Trem2*, *Mafk*, *Mmp7*) were also validated in RNA-seq datasets of patients with cirrhosis and portal hypertension. Finally, deconvolution analysis identified major cell types and suggested an association of macrophage and portal hepatocyte signatures with portal hypertension and fibrosis area in both models.

## Introduction

Liver fibrogenesis is a complex process characterized by functional alterations in multiple cell types and excessive extracellular matrix (ECM) turnover in response to ongoing hepatic injury^1^, which can ultimately result in cirrhosis and portal hypertension (PH)^2^. Despite significant progress in understanding the pathomechanisms contributing to liver fibrosis^3^, the molecular drivers of PH and fibrosis regression still remain elusive. Investigation of hepatic gene expression patterns (i.e., transcriptomic signatures) associated with PH severity and liver fibrosis regression will not only enhance our comprehension of liver disease but also holds promise for the development of novel therapeutics for patients with liver cirrhosis and PH.

Persisting hepatic necroinflammation^4^, vascular remodelling characterized by capillarization of liver sinusoidal endothelial cells (LSEC)^5^, macrophage activation^6^, and transdifferentiation of hepatic stellate cells (HSC) to myofibroblasts^7^ are hallmarks of liver fibrosis. Although it is well-known that the removal of the primary etiologic factor of liver disease enables patients to achieve fibrosis regression^8^ and amelioration of PH^9^, resulting in the regeneration of hepatic function, there has been no systematic analysis of the of hepatic gene expression dynamics associated with these processes in a controlled setting.

Nevertheless, the dynamics of hepatic gene expression associated with liver fibrosis regression and PH severity have yet to be systematically analysed in a controlled setting, such as in widely used mouse models of carbon tetrachloride (CCl_4_)- or thioacetamide (TAA)-induced liver disease.

Mechanistic insights on the molecular drivers of portal hypertension (PH) suggest that endothelial dysfunction^10^, pathologic angiogenesis^11^, and abnormal HSC-LSEC crosstalk^12^ are critical factors. However, obtaining high-quality human liver biopsy material for research remains challenging, thus limiting the application of high-throughput technologies in studying the dynamics of liver disease progression and regression in human patients in an unbiased fashion.

The use of animal models provides a unique setting to investigate main molecular pathways involved in liver fibrosis regression and PH severity. Our study design included spontaneous regression after removing the inducing agent for up to two weeks, allowing us to identify molecular markers linked to the disease dynamics, correlate them to key liver disease surrogates, and discover related regulatory transcriptional factors (TFs). To this end, we generated and analysed transcriptome profiles in two well-established mouse models of advanced fibrosis induced by CCl_4_ and TAA^13, 14^. We characterized the regulons of these TFs and proposed pathways linking them to fibrosis regression and PH severity. Hub genes were validated using publicly available human RNA-seq datasets. Finally, shifts in deconvoluted single-cell signatures suggested a critical contribution of macrophages, non-parenchymal cells, and hepatocytes to liver fibrosis and portal hypertension.

## Results

### Disease characteristics of the parallel fibrosis progression and regression in animal models

Key liver disease parameters were assessed at the same timepoint when liver samples for RNA sequencing (RNA-seq) were harvested, specifically after 12 weeks of vehicle administration for healthy control (HC) or 12 weeks of toxin exposure inducing peak fibrosis (cirrhosis, CIR), and then after one week (R1) or for two weeks (R2) of spontaneous fibrosis regression (Figure 1a, b). Portal pressure (PP), liver fibrosis (collagen proportionate area, CPA), and hepatic injury (reflected by alanine (ALT) and aspartate (AST) aminotransferase blood levels) were significantly increased in CIR animals (i.e., at peak induction) of both models induced by carbon tetrachloride-induced (CCl_4_) or thioacetamide (TAA), and subsequently decreased during R1 and R2 regression (Figure 1c). Fibrosis induction resulted in a significant increase in PP in induced animals. We observed CIR_TAA_ median level of 7.47 mmHg, and CIR_CCl4_ animals reached the highest values with a median of 10.41 mmHg (Table 1). Median CPA levels were below 1% in control animals and were increased 5-fold (CIR_TAA_: 3.92%) and 20-fold (CIR_CCl4_: 16.81%) in diseased groups (Figure 1c, d). Both ALT and AST levels were elevated at peak fibrosis, however, at lower intensities in TAA than in CCl_4_. Both the CPA and portal hypertension decreased during the regression phase. The decrease in PP was not significant at R1 in the TAA model, improving only in R2_TAA_. The levels of transaminases (ALT, AST) significantly decreased in both models already at the R1 timepoint. The course of these key liver disease readouts indicates that the models and timepoints are well-suited to study gene regulation at peak fibrosis and portal hypertension and during liver disease regression, which was subsequently confirmed by unsupervised principle component analyses.

**Figure 1.**
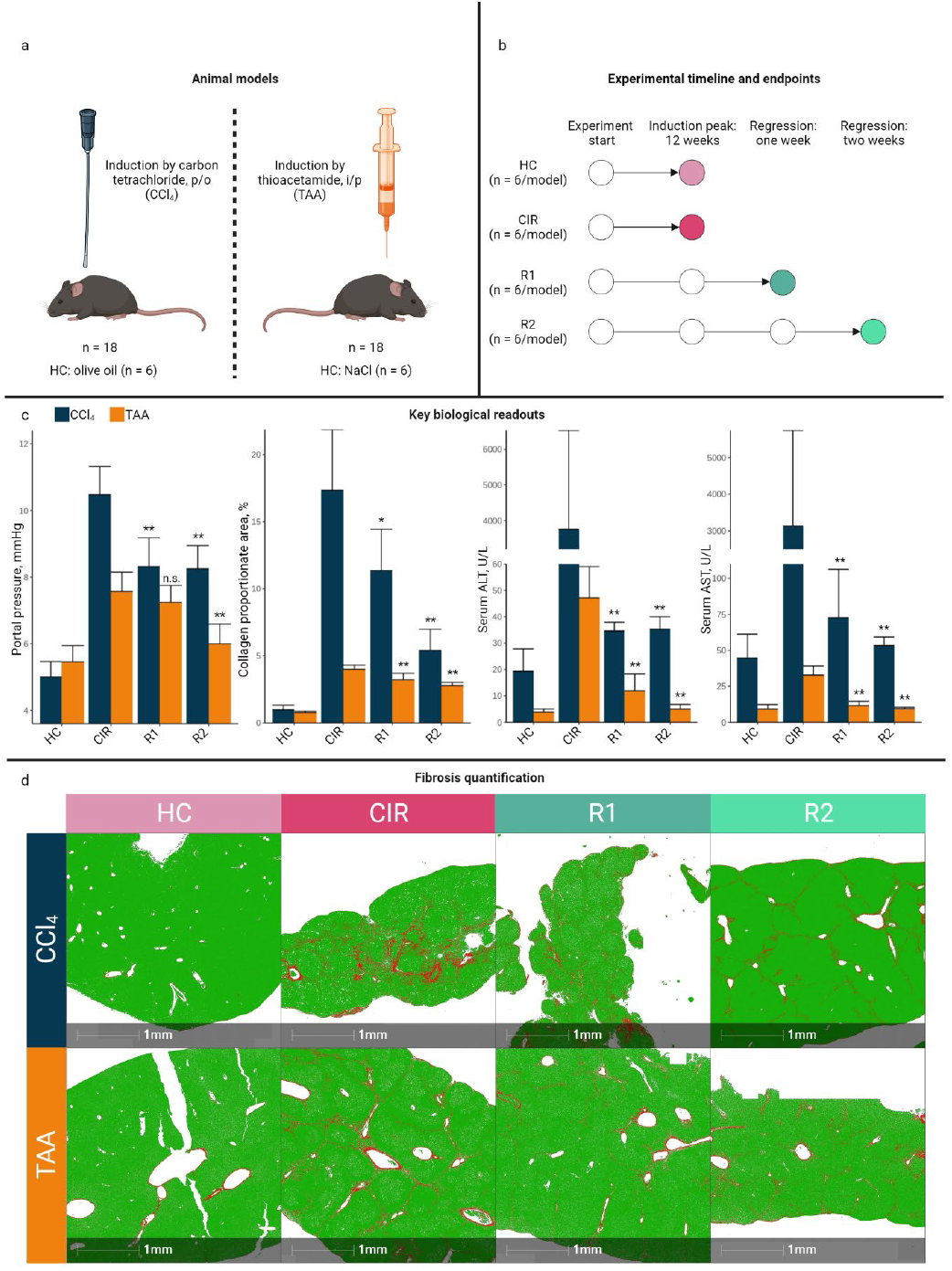
Experimental design and key liver disease readouts. **(a)** Two different murine fibrosis models were used to study their hepatic transcriptome during peak fibrosis and fibrosis regression. **(b)** Experimental timelines are shown. Portal pressure was measured, blood was sampled, and liver tissue was harvested at the respective endpoints. **(c)** Comparison of key liver disease readouts in the two models. Results of the Wilcoxon rank-sum test for the differences in the respective parameters after one (R1) and two (R2) weeks of regression to the peak (CIR) time point are indicated (n.s. = non-significant; * p<0.05; ** p<0.01; n = 6 per group of each model). **(d)** Illustration of representative Picrosirius Red staining for quantification of collagen proportionate area are shown. Red = collagen, green = fast-green tissue counterstain.

**Table 1.**
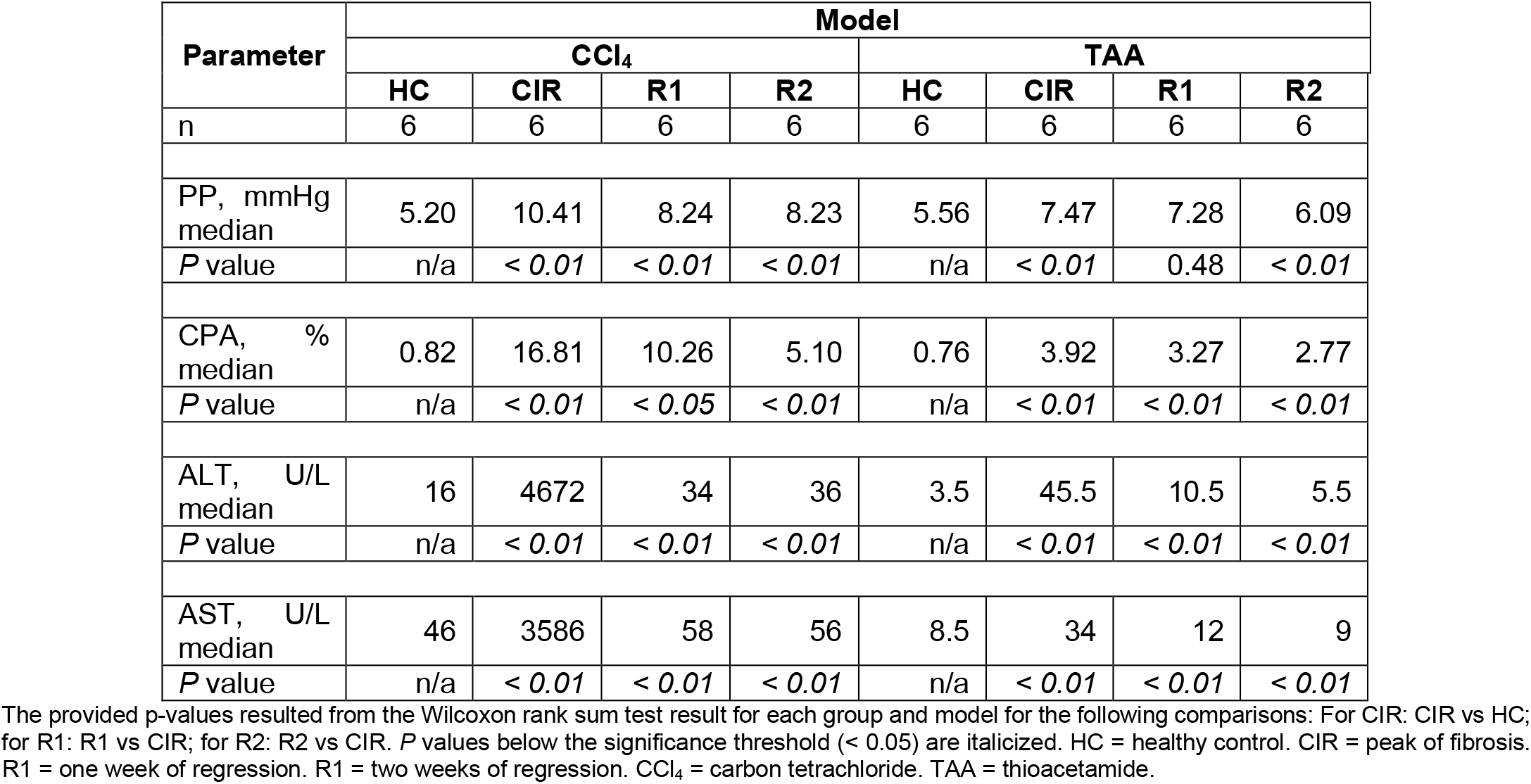
Characteristics of the animal cohort.

### Immune and matrix remodelling-related genes are driving transcriptomic variance in fibrosis

We used principal component analysis (PCA) to assess group similarity and identify genes defining these groups. Whole transcriptomes of the sequenced animal livers were provided as an input, and the first three components adequately distinguished CIR and HC while grouping R1+R2 together (Supplementary Figure 2-1). Principal components (PC) 1 and 2 together explained 52.3% of the variation of the dataset (Figure 2a). Still, some CIR_CCl4_ animals appeared to be grouped with R1+R2, suggesting that - even in a controlled environment – some heterogeneity in the transcriptomes of peak fibrosis in the CIR_CCl4_ model can be expected. Following PCA, the top 10% of the genes explaining variance in components 1 and 2 were retained (n = 22) (Figure 2b). We found matrix metalloproteinase-7 (*Mmp7*) – that is reportedly involved in ECM remodelling and cell adhesion^15^ – to be upregulated at peak fibrogenesis and with maintained expression throughout the regression phases, except in some R1+R2_TAA_ animals (Figure 2b). In the CIR cluster, genes *Cd63*, *Ly6d*, and *Cdkn1a,* involved in immune cell activation, were all associated with peak fibrosis. The stearoyl-coenzyme A desaturase 4 (*Scd4*), Abhydrolase Domain Containing 1 (*Abhd2*), and *Mia2* genes were overexpressed in CIR_TAA_ but not to the same magnitude in CIR_CCl4_. In contrast, Indolethylamine N-Methyltransferase (*Inmt*), responsible for xenobiotic degradation, showed reduced expression in CIR_TAA_ but not CIR_CCl4_, returning to control levels already after one week of regression. Two solute carrier genes, *Slco1a1* and *Slc1a2*, were strongly downregulated by fibrosis induction and not completely restored after two weeks for both models: *Slco1a1* is a transporter of organic anions shown to be dramatically downregulated in ethanol-induced liver damage^16^. *Slc1a2* is involved in glutamate transport, which was previously reported to be downregulated in CCl_4_ cirrhosis model and identified as a hub gene in fibrosis^17^.

**Figure 2.**
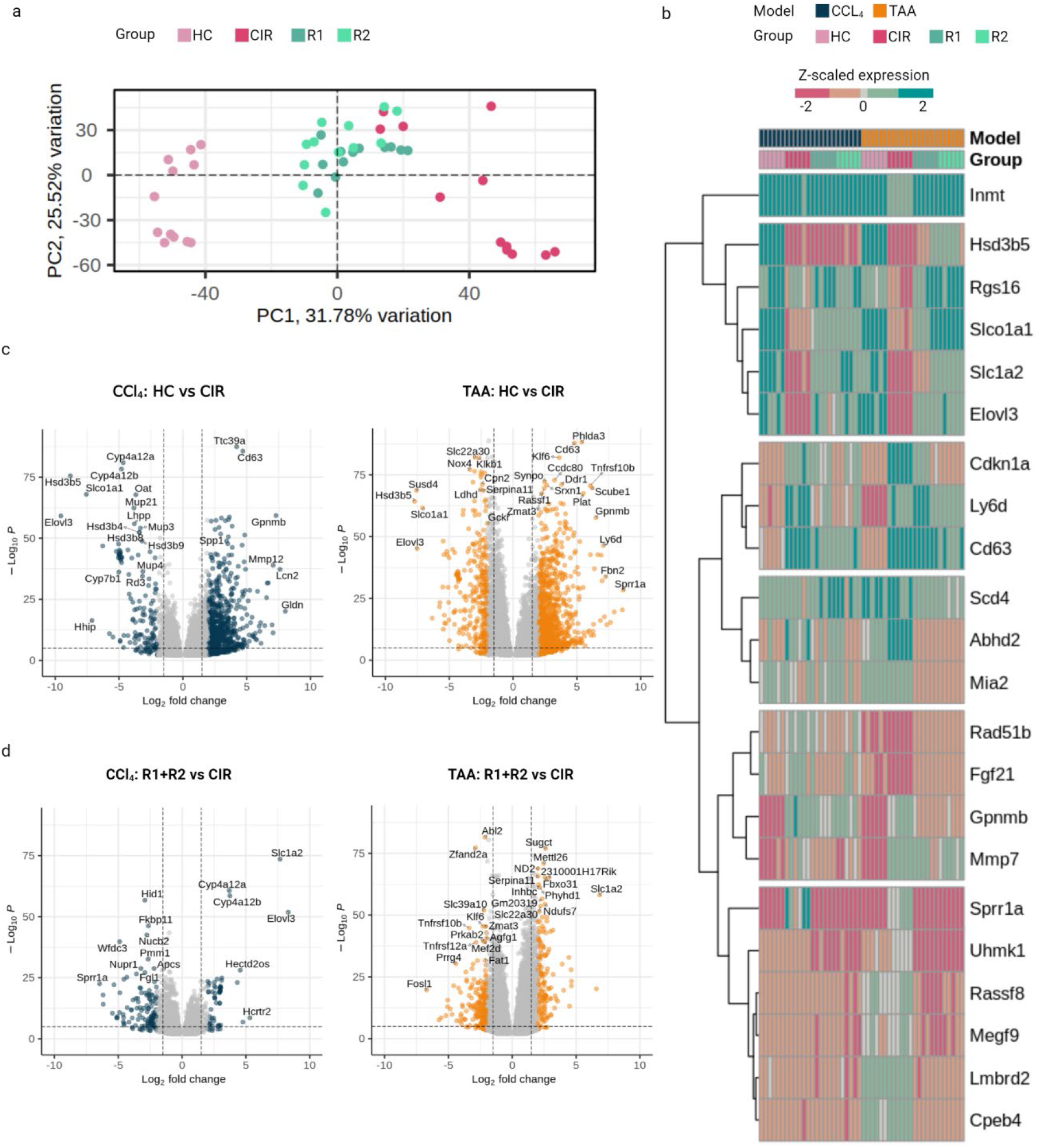
Transcriptomic characterization of genes associated with peak fibrosis and regressive fibrosis. **(a)** PCA projection demonstrates a clear separation of HC and mostly of CIR, while the R1 and R2 groups cluster together. **(b)** Genes with 10% highest loading score for principal components 1 (PC1) and 2 (PC2). Unsupervised gene clustering denotes similar expression patterns. **(c)** Volcano plots for most differentially expressed genes between animals at peak fibrosis induction (CIR) and healthy control animals (HC) are shown separately for the CCl_4_ and TAA models. **(d)** Volcano plots for most differentially expressed genes between animals with regressive cirrhosis (combined R1+R2) vs. peak fibrosis induction (CIR) are shown separately for both models. Thresholds: log2FC > 1.5, padjusted < 0.01. PC = principal component.

Consequently, we performed differential expression analysis to characterize each model independently. It resulted in 1547 significantly differentially expressed (DE) genes between CIR_CCl4_ and HC_CCl4_, and 2614 genes between CIR_TAA_ and HC_TAA_ (Supplementary Data 1, 2). In these groups, upregulation of extracellular matrix-related genes in both models was observed, with *Cd63* and *Ccdc80* overlapping among top-upregulated genes (Figure 2c). Downregulation in both cases included cytochrome family genes (prominently *Cyp4a12a*, *Cyp4a12b*, *Cyp2j5*), and solute carriers with a less well-defined role in liver fibrosis (*Slc22a30*, *Slc22a28*). Notably, *Cyp2j5*, involved in arachidonic acid metabolism, was not previously discussed in the fibrogenesis context but was reported as downregulated in a NAFLD mouse model^18^.

Following these findings, we assessed extracellular matrixrelated markers among DE genes. In both CIR groups, collagens *Col5a2*, *Col1a1,* and Fibrillin 1 (*Fbn1*) were highly up-regulated – as previously reported for both early and advanced fibrosis ^19, 20^. Fibrillin 2 (*Fbn2)* also was among these markers, but its function is less understood. Notably, *Fbn2* was downregulated in acute CCl_4_ injury^21^, but, in line with our findings, it was upregulated in 6-week CCl_4_ administration in *Pxdn* deficient mice^22^. *Mmp12*, a macrophage-derived enzyme involved in the degradation of elastic fibers, and IL13, mediating hepatic stellate cell (HSC) activation^23^, were upregulated in both models. CIR_CCl4_ had a more prominent upregulation of *Timp1,* hinting at more advanced liver injury^24^. Upregulation of *Scube1* might be of compensatory or protective nature since its downregulation has been linked to endothelial damage severity in pulmonary hypertension and fibrosis^25, 26^. Top-downregulated genes included *Capn8* and *Col27a1*, indicating degradation of the homeostatic matrix, and *Klkb1*, providing a functional link to coagulation (Supplementary Figure 2-3a, b).

Even though both R1_CCl4_ and R1_TAA_ had already a large set of DE genes compared to the fibrosis peak, there were no DE genes between R2 and R1 for the CCl_4_ model. For TAA, one gene was significantly upregulated in R2 compared to R1 (*Slc1a2*), and two were downregulated (*Cdhr2*, *Dpf1*). *Dpf1* negatively regulates transcription, and *Cdhr2* is involved in contact inhibition and regulation of cell proliferation, suggesting that their functions in fibrosis are of protective nature that are engaged in long-term regression. Among other matrix-related genes, *Vtn*, a target of Mmp2, was strongly upregulated in R1+R2_TAA_ (Supplementary Figure 2-3c, d). In both models’ R1+R2, a substantial upregulation of cytochrome subunits was observed with a total of 17 genes in TAA and six overlapping in CCl_4_. Most of them were involved in phase I of the P450 pathway, primary bile acid synthesis, retinol, and steroid metabolism. *Cyp1a1*, involved in xenobiotic metabolism, was slightly downregulated in both models’ regression. *Timp1* (CCl4), *Itga3*, *Itga6*, and *Itgav* (TAA) were significantly suppressed compared with the fibrosis peak. *Col5a3*, however, still showed increased expression in R1+R2_TAA_, suggesting that HSC activity was maintained two weeks after the removal of the injury agent. The *Slc1a2* was upregulated in both models after being suppressed at peak fibrosis (Figure 2d, Supplementary Data 3, 4).

Together, these results indicate that both models display an upregulation of classical markers of liver fibrosis, which are then partially again downregulated in disease regression. Other less-studied DE genes also detected, but their role in liver fibrosis should be further clarified. Considering the strong grouping of the liver transcriptomes of animals during fibrosis regression on PCA and the DE results, we kept R1+R2 (R) merged for the following analyses.

### CCl_4_ and TAA induce model-specific signatures and distinct recovery patterns

We compared matching groups in two models to identify further functional alterations introduced by CCl_4_ and TAA. From DE analysis, 30% (n = 959) of genes overlapped when comparing CIR_CCl4_ and CIR_TAA_ to their respective HC groups (Figure 3a). The upregulation overlap included pathways involved in focal adhesion, cell cycle control, prostaglandin signalling, fibrosis, and macrophage regulation (Figure 3b). The most robustly downregulated pathway was PPAR signalling, followed by the P450 pathway, steroid, fatty acid, and amino acid metabolism. Our findings aligned with the current understanding of the PPAR role in advanced chronic liver disease, as pan-PPAR agonists have shown promising results on fibrosis resolution both in an experimental studies and in phase 2b clinical trials^27, 28^.

**Figure 3.**
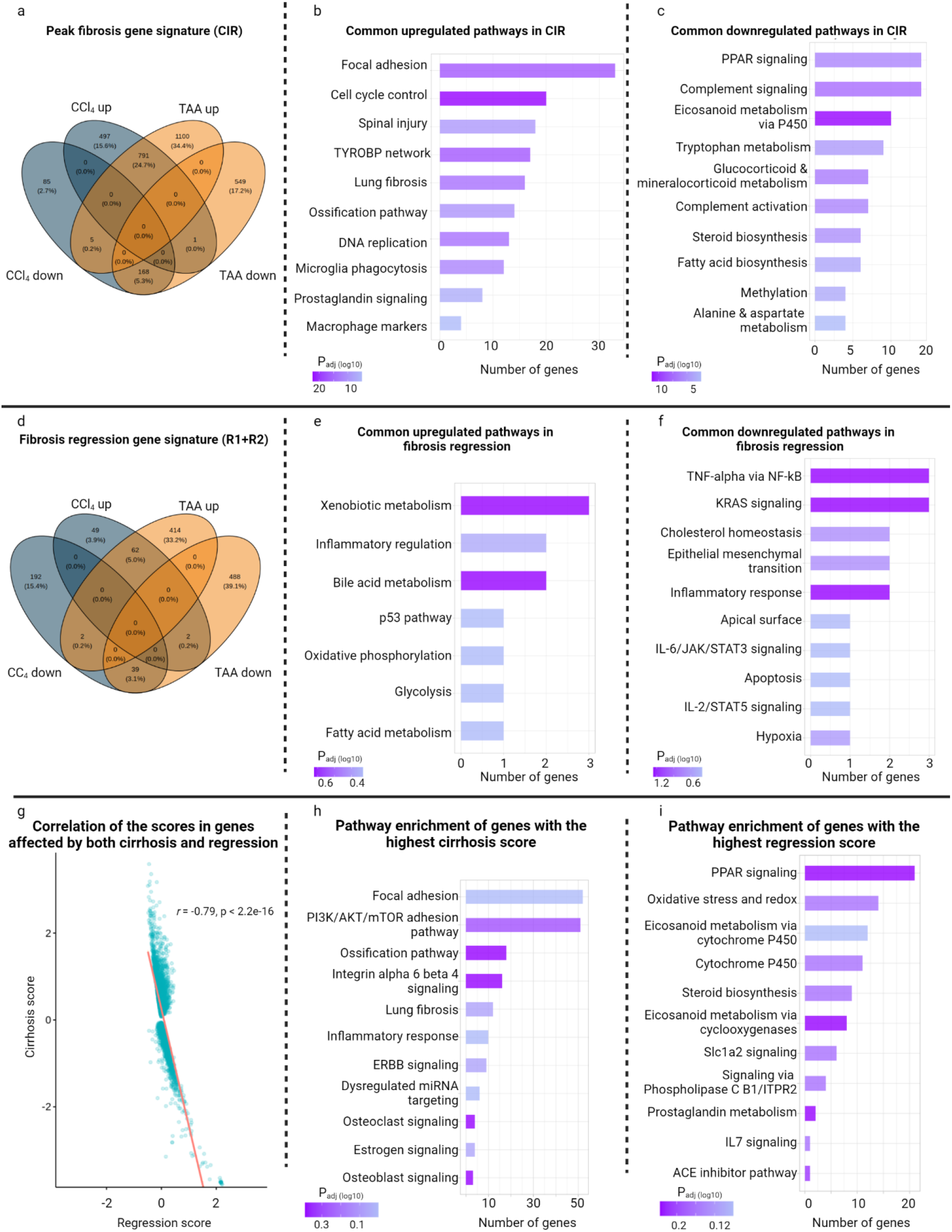
Functional annotation of genes involved in CCl4 and TAA fibrosis progression and regression. **(a)** Venn diagram representing overlaps in up- and downregulated genes (number and proportion of genes are shown) associated with fibrosis induction in the two fibrosis models. The functional analysis displays molecular signaling pathways for both CCl4/TAA models that are **(b)** up- or **(c)** downregulated in peak fibrosis induction (CIR). **(d)** Venn diagram displaying overlaps in genes (number and proportion of genes are shown) linked to fibrosis regression in the two models. **(e)** Functional upregulation and **(f)** downregulation of molecular signaling pathways for both CCl4/TAA models during fibrosis regression. A hypergeometric test with MSigDB annotation was applied for the genes overlapped in regression. **(g)** The scores in genes perturbed by both conditions demonstrate a strong negative correlation (Pearson’s *r* is shown). The scores were scaled for visualization. **(h)** Signaling pathways from genes with the 20% highest score for CIR are shown. **(i)** Pathways from genes with the 20% highest regression score (CIR vs. R1+R2) are shown.

Model-unique genes suggested that CIR_CCl4_ had a more substantial downregulation of the PPAR pathway than CIR_TAA_, and a more prominent signature of metabolic inhibition (Supplementary Figure 3-1a, b). Notably, cholesterol biosynthesis was preserved in CIR_CCl4_. The delta-Notch pathway, involved in portal fibrosis and tissue remodelling in cirrhosis^29^, was inversely regulated: it was upregulated in TAA peak induction but downregulated in CCl_4_. In differential testing of the CIR groups, we found that *Pdzk1*, *Ccl9,* and *Ppp1r42* were among the highest expressed markers of CIR_CCl4_, while *Ets2*, *Slc39a10*, and *Fem1b* were associated with CIR_TAA_ (Supplementary Figure 3-2a, Supplementary Data 5).

The regression overlap of both models included 8.1% genes (n = 101) (Figure 3a). The upregulation of xenobiotic metabolism, inflammatory regulation, and bile acid metabolism was observed together with the downregulation of classical proinflammatory pathways (NF-kB, KRAS, IL6-STAT3) (Figure 3b, c). Unlike TAA, R_CCl4_ showed a significant upregulation of the WNT signalling. A restoration of PPAR and metabolic signalling was observed in CCl_4_, while for TAA, the highest pathways were related to oxidative phosphorylation (Supplementary Figure 3-1c, d). We detected significant upregulation of *Ppargc1a* (coactivator of *Pparg*) and PPAR-beta/delta genes and their downstream mediators *Fabp1*, *Acaa1a*, *Ilk*, *Scd1*, and related cytochromes (Supplementary Data 3) in fibrosis regression.

Top R_TAA_-specific genes included *Thbs4* and *Dmbt1* (Supplementary Figure 3-2b, c). The latter was previously reported as highly expressed in liver regeneration in rats^30^. *Cd163* and *Hhip* were specific to R_CCl4_. In previous studies, Hhip was shown as an HSC-derived gene and a member of the Sonic hedgehog (Shh) pathway that is upregulated when HSC activation is prevented^31^ (Supplementary Data 6).

These functional overlaps revealed that both models express preserved fibrogenesis pathways, followed by the downregulation of inflammatory pathways in regression and upregulating of metabolic and oxidative phosphorylation pathways (Supplementary Data 7). The identified model-specific differences in delta-Notch, PPAR, Wnt, and Sonic hedgehog pathways should be considered when testing therapeutic candidates.

As an alternative strategy to differential gene expression testing, we investigated how genes are perturbed by phenotype change toward fibrosis or regression. Both models were merged for these experiments, and expression in the HC or CIR group was considered a baseline to calculate cirrhosis and regression scores (see “Methods”). These scores represented each specific gene’s expression alteration towards CIR or R accordingly. We found that most of the genes were altered by cirrhosis towards upregulation and in regression towards downregulation (Figure 3g) – indicating injury-driven gene upregulation in peak fibrosis. In genes impacted by both perturbations, a strong negative correlation between the scores was detected (Supplementary Figure 3-3).

Next, functional analysis was performed with genes ranked as the 20% highest absolute scores. Genes with the highest cirrhosis score showed involvement in adhesion, fibrosis, and integrin-related signalling (Figure 3h). Similar to DE analysis findings, pathways like PPAR, P450, steroid biosynthesis, and oxidative phosphorylation were upregulated among genes with the highest regression score (Figure 3i). The above-mentioned pathways were mostly reversely overlapping in the lowest 20%-scored genes (Supplementary Figure 3-4).

### Pseudotemporal dynamics reveal gene trajectories in disease progression and regression

By jointly leveraging the gene expression data from the models, we endeavoured to gain insights into the evolution of disease and recovery. The aim of this analysis was to identify genes perturbed by these processes and the direction of changes over experimental endpoints – to which we refer as pseudotemporal dynamics. We defined all our study groups as pseudotemporal points. We investigated how the expression of the same genes changes between the non-diseased state (i.e., the HC group), the peak of induction (CIR), and one to two weeks without the damaging factor (R1+R2). The genes changing in the same direction were then clustered, and the following patterns were observed (Supplementary Figure 4-1):

(Cluster 1) Upregulated in fibrosis, then downregulated in regression (1273 genes);

(Cluster 2) Downregulated in fibrosis, upregulated in regression (n = 1225);

(Cluster 3) Upregulated in fibrosis, upregulated in regression (n = 261);

(Cluster 4) Downregulated in fibrosis, downregulated in regression (n = 241).

On the pathway level, clusters 1 and 2 were mostly well in line with our findings from DE genes for CIR vs. HC and R vs. CIR accordingly, as well as with top-ranked genes for perturbation signatures. The most prominent pathways in cluster 1 were cell adhesion, followed by progenitor cell signalling, and profibrotic pathways, including TGFβ and Dmp1 signalling. Cluster 2, in turn, had PPAR as the most prioritized pathway, followed by amino acid and cholesterol metabolism, oxidative phosphorylation and the P450 pathway (Figure 4a, b).

**Figure 4.**
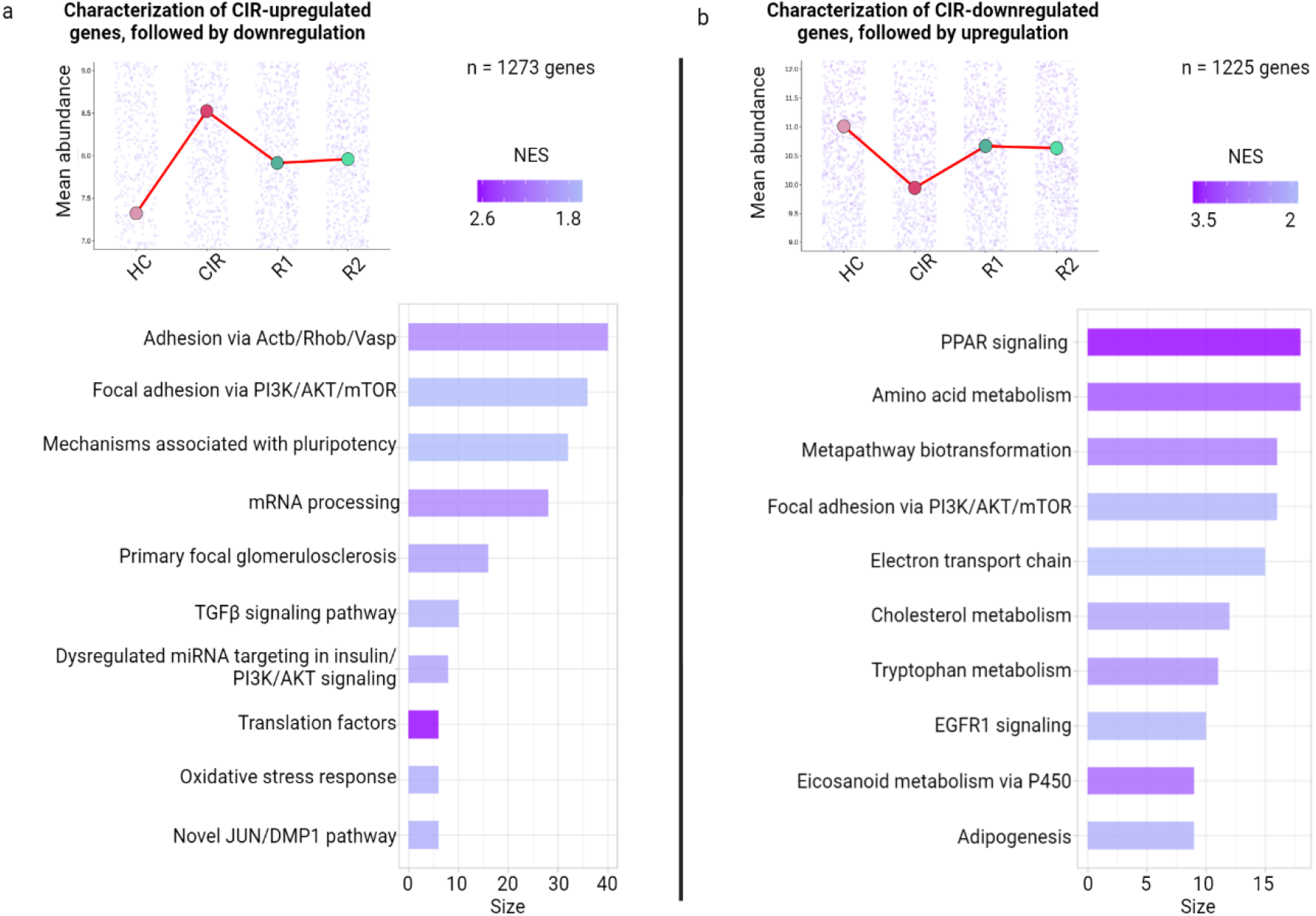
Dynamic signature patterns following liver disease course using pseudotemporal analysis. **(a)** Cluster 1: pathway analysis of genes upregulated at peak fibrosis followed by subsequent downregulated (vs. CIR) and approximation to baseline (HC) levels in regression. **(b)** Cluster 2: pathways analysis of genes downregulated at peak fibrosis followed by subsequent upregulation (vs. CIR) and approximation to baseline (HC) levels in regression. NES = normalized enrichment score.

Cluster 3’s pathways were similar to those described in CIR and cluster 1, with cell adhesion via *Itga8* and laminins playing the most prominent role. Another finding was persisting upregulation of matrix metalloproteinases, mainly *Timp3*, *Timp2*, and *Mmp2* – which suggests their role in sustained matrix remodelling even in the regression stages (Supplementary Figure 4-2a).

Cluster 4 was characterized by a slight downregulation in both fibrosis and regression. Its genes were not strongly grouped in pathways, with highest (yet not significant) ranked ones belonging to proteasome degradation and oxidative damage. The macrophage marker *Cd163* was found among cluster genes, which is known to be downregulated in the proinflammatory microenvironment. Additional findings included solute carriers transporting nucleic bases, prostaglandins, and zinc (*Slc35e3*, *Slco2a1*, *Slc39a4*), and *Scarb2*, one of the scavenger receptors on the LSEC membrane (Supplementary Figure 4-2b).

Cluster 2 had the most extensive overlap with DE genes for all groups (n = 473 genes), followed by cluster 1 (n = 316). The most considerable overlap of cluster 3 was with genes DE for both CIR (n = 95), and cluster 4 contained most DE genes of R_TAA_ (Supplementary Figure 4-3). These response patterns emphasized the plethora of molecular alterations brought by fibrogenesis and its disease-associated dynamics (Supplementary Data 8-11). Aside from impacting extracellular matrix composition, it also suggests the involvement of hepatic non-parenchymal cells, reflected by their hallmark genes.

### Discovery of transcriptional biomarkers via co-expression and targeted learning

Two independent methods were applied to narrow down the list of DE genes, and biomarker candidates were identified in the overlap of their results. We selected the first one, WGCNA, as an established unsupervised method for finding genes co-expressing with external parameters. The second one, TMLE, is a machine learning-based approach to estimating gene expression impact on selected liver disease parameters while controlling for other confounders. In both cases, we aimed to identify markers related to the key biological liver disease severity readouts PP and CPA as continuous values and using cirrhosis and regression as categorical parameters. This biomarkers search was performed model-agnostic and using merged regression groups.

Weighted co-expression network analysis (WGCNA) discovered gene modules strongly linked to our external readouts (Figure 5a). PP, CPA, and cirrhosis shared the correlation direction of such significant modules; the shared injury signature was also for ALT and AST gene modules.

**Figure 5.**
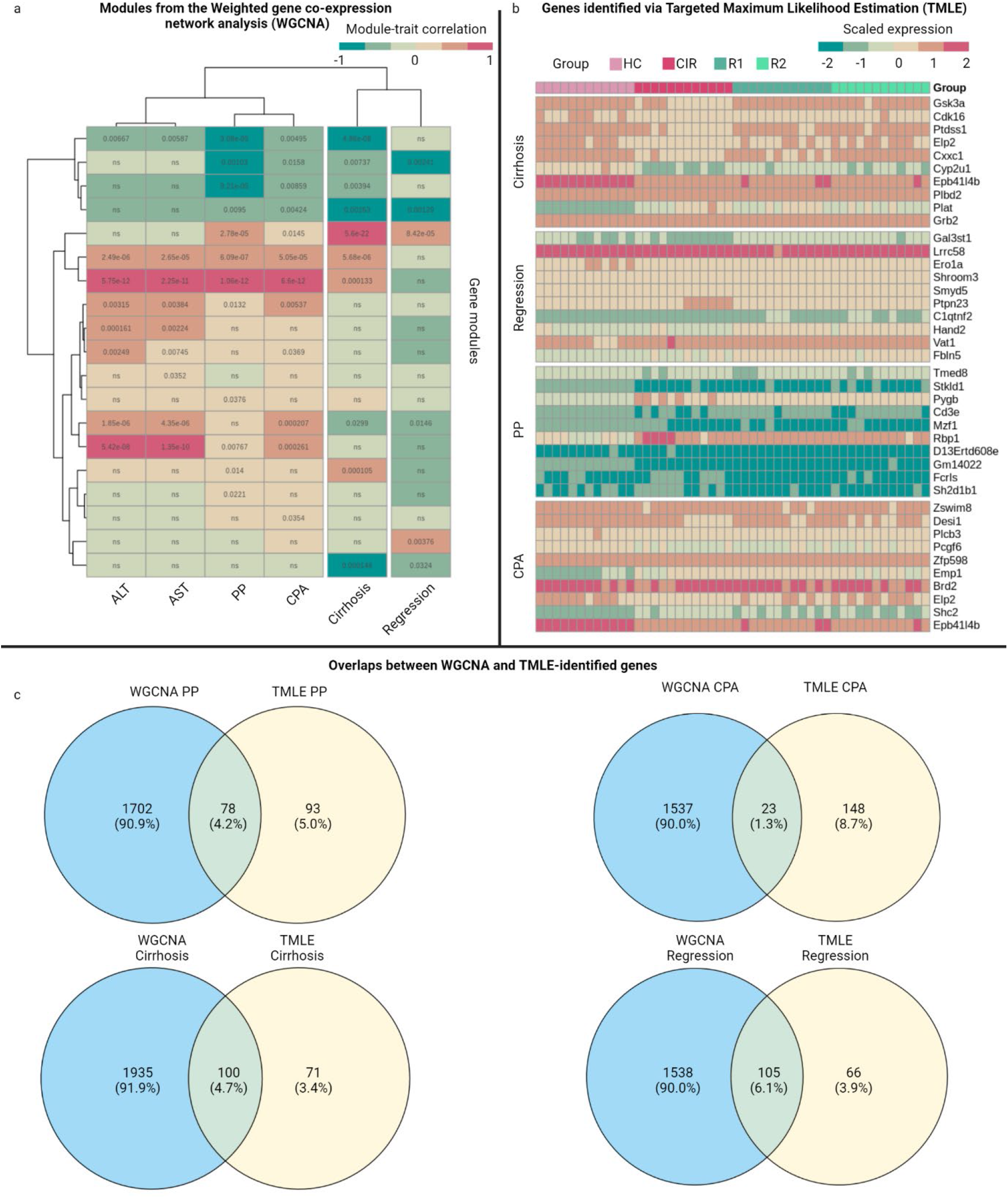
Linking gene signatures to key surrogate parameters of hallmark biological readouts. **(a)** Gene modules, identified via unsupervised weighted co-expression network analysis (WGCNA), correlating with key biological readouts in the merged dataset. **(b)** Supervised clustering of top 40 markers associated with Targeted Minimum Loss-Based Estimation (TMLE). **(c)** prioritization via overlaps between TMLE markers for each biological readout and respective WGCNA module. PP = portal pressure. CPA = collagen proportionate area.

The key pathways in the gene set co-expressed with regression were Myc targets or unfolded protein response (Supplementary Figure 5-1). The top markers included genes involved in vascular signalling (e.g., *Vegfa, Angptl2*) and matrix-related genes (e.g., *Adam33*, *Adamtsl2*, *Igfbp7*) (Supplementary Data 12).

The highest-scored genes associated with portal pressure were related to pathways involved in epithelial-mesenchymal transition, coagulation, and xenobiotic metabolism. It also contained HSC activation state markers such as *Timp3*, *Wnt5a*, *Vim*, and *Pdgfb*. Identification of both HSC and sinusoidal markers (*Pecam1*, *Vwf, Cd9*) among highly significant genes for PP in these animal models supports previous reports on the involvement of the vascular microenvironment in the development of hemodynamic alterations^32–34^. Furthermore, both cirrhosis and CPA-linked genes comprised a vast number of matrix turnover genes, as well as angiogenesis genes (e.g*., Adam8, Itgav, Vegfa, Pdgfa, Vcan*). In addition to mesenchymal-epithelial transition, angiogenesis was among the high-scored pathways for both gene sets.

ALT and AST share a common signature, including E2F targets, known as a pro-apoptotic pathway in the liver, xenobiotic metabolism, adipogenesis, and bile acid metabolism. Among the highest-ranked genes were Prtn3, a monocyte marker; Vsig8, a regulator of T-cell activation; *S100a3*, a calcium-binding protein regulating gene expression in hepatocytes and a potential treatment target in hepatocellular carcinoma^35^.

Targeted Minimum Loss-Based Estimation (TMLE) identified that Aldo-keto reductase family 1 member C18 (*Akr1c18)* and Nucleobindin 2 (*Nucb2*) were linked to higher PP (Supplementary Figure 5-2a). *Cyp2u1*, on the opposite, was downregulated in CIR animals with the highest level of portal pressure (Figure 5b). A cluster of genes involved in matrix turnover showed an association with CPA and cirrhosis: *Spire2*, *Pak6*, *Tinag*, and HSC-annotated markers *Fhl2*, *Tinag*, *Tgfb2*, and *Shc2* (Supplementary Figure 5-2b). In line with WGCNA, the TMLE prioritized *Igfbp7*, linked to regression together with *Cd74 (Supplementary Data 13)*.

Using overlaps of these two methods, we identified biomarker candidates for PP (n = 78 genes), CPA (n = 23), cirrhosis (n = 100), and regression (n = 105) (Figure 5c, Supplementary Data 14). The overlapping genes for PP were classified as scavenger receptors, enzymes, and members of RAS, PI3K, and FYN pathways (Supplementary Figure 5-3a). The CPA-linked markers included genes coding proteoglycans, elastic fiber, scavenger receptors, and members of the RUNX2 pathway (Supplementary Figure 5-3b). The set of genes overlapping in cirrhosis consisted of ECM turnover-related genes, chemokine receptors, elastic fiber, and matrix metalloproteinases (Supplementary Figure 5-3c). Notably, regression genes were also involved in the turnover of elastic fibers; additionally, members of Tie2, PTK2, and apoptotic pathways were found (Supplementary Figure 5-3d).

### Transcriptional regulators in fibrosis progression and regression and their signalling niches

To identify transcription factors (TFs) involved in the perturbation of the identified transcriptomic signatures of fibrosis progression and regression, we used differential co-expression of the annotated mouse TFs based on their known target genes (see “Methods”).

A total of 58 TFs were identified as significantly involved in our dataset (Supplementary Data 15). We examined their expression and discovered that three of them (*Hmga1*, *Tcf4*, *Mafk*) are significantly upregulated in peak fibrosis (P < 0.01), while significant downregulation of *Klf15*, *Ppara*, *Xbp1*, *Creb3l3*, *Foxo4*, *Nfe2l1*, *Nr1d1*, *Nr1i2m*, and *Junb* was detected in CIR. When exploring potential functional links, *Creb3l3* seems to be implicated in anti-inflammatory control since its knockout aggravated liver fibrosis in mouse models^36^. Notably, the downregulation of Foxo4 was previously described in the context of proinflammatory and profibrogenic microenvironment^37^. The identified TFs belonging to the nuclear receptor family included *Nr1i2*, *Nr1h2*, *Nr1d1*, *Rxrb*, and *Ppara*, and their functions were previously implicated in liver disease^38^.

A protein-protein interaction network was constructed using significant TFs and all DE genes between the study groups to perform their functional annotation (Supplementary Figure 6a). To have a broad perspective of their downstream, we applied the “guilt by association” principle^39^ and identified genes directly connected to the TFs, or their 1^st^degree neighbours. Functional annotation of the resulted network showed that the identified TFs are involved in the regulation of several immune pathways, with NF-kB and TGFβ among the most prioritized ones (Supplementary Figure 6b). Other findings included Wnt/β-catenin, Notch, Hedgehog, adipogenesis and junction pathways, thus complementing our previous findings from the DE analysis. Next, we aimed to narrow down further the list of significant TFs relevant for regulating the biomarker candidates identified within WGCNA-TMLE overlaps. Their associations were scored by the expression changes in the biomarker genes, and the weakest links were discarded. It allowed us to select 22 TFs impacting levels of PP and CPA, cirrhosis and regression (Supplementary Data 16). In the following experiment, we integrate these findings via network analysis.

### Analysis of multi-layer network results in prioritized biomarkers and their functional links

In previous experiments, sets of biomarker gene candidates and their transcriptional regulators were identified. We assumed that analysis of their biological network could allow us to further select the most informative genes for validation. For this analysis, a three-layer network was constructed: a gene regulatory layer (n = 22 genes) consisting of TFs; a gene co-expression layer (n = 299), including biomarkers, overlapped from the described discovery approaches; a protein-protein interaction layer (n = 751), including database-mined downstream and upstream targets for the biomarker candidates (Figure 6a).

**Figure 6.**
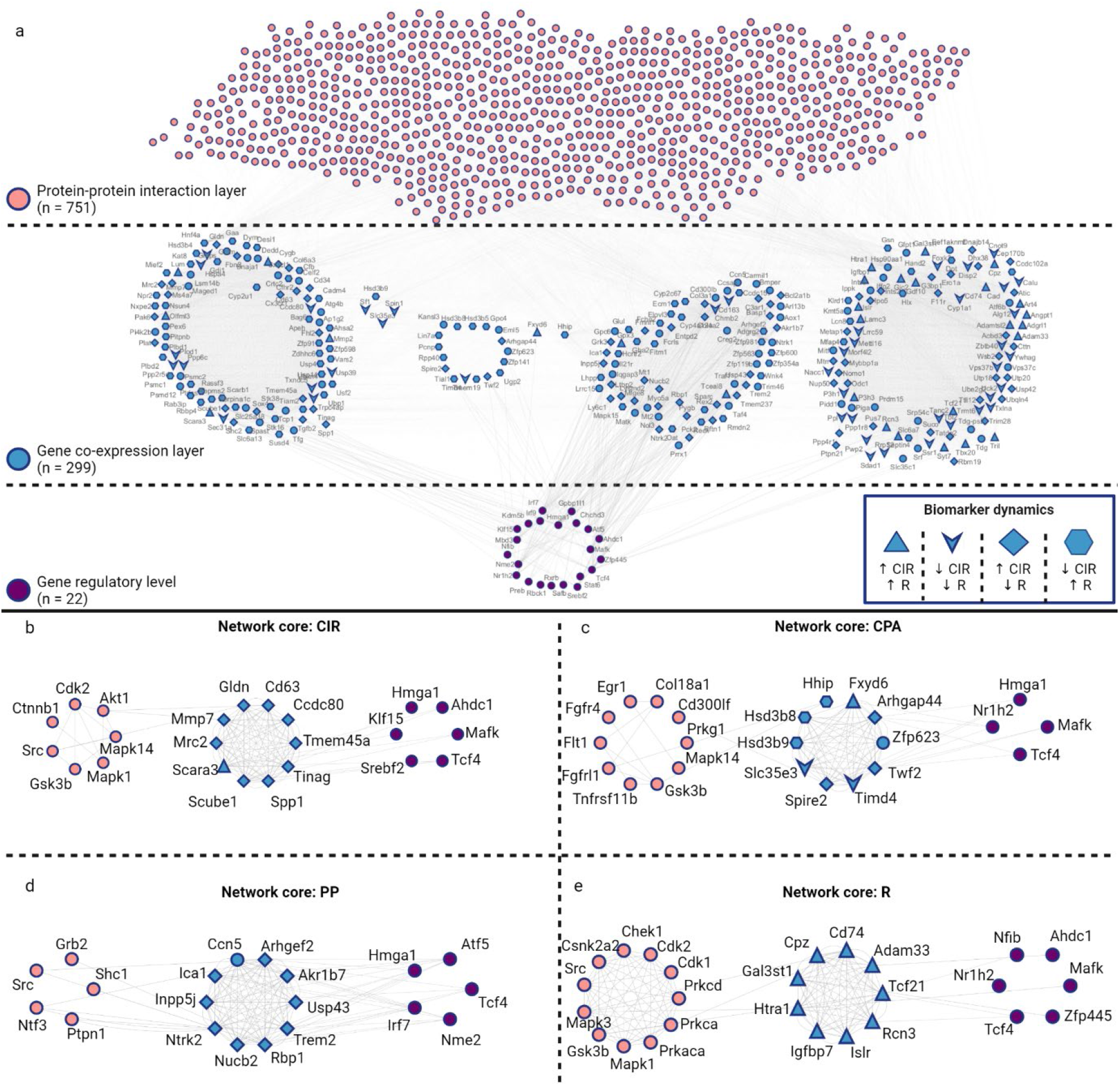
Network analysis prioritizes biomarkers and shows their functional links to liver disease. **(a)** Multilayer network constructed from selected transcriptional factors (purple, bottom layer), co-expressed biomarkers (blue, middle layer), and their protein-protein interactions (orange, top layer). **(b-e)** Biomarkers were scored to identify network cores with the feature’s strongest associations. Node shapes indicate gene dynamics between cirrhotic and regression groups. CIR = cirrhosis, CPA = collagen proportionate area, PP = portal pressure, R = fibrosis regression.

For prioritization, we employed biology-based scores (perturbation scores that were calculated earlier), several network-based scores, and a machine-learning algorithm (see “Methods”). Using combinations of these scores, the network cores (up to n = 25 genes) were identified for each disease “phenotype” category: cirrhosis, CPA, PP, and regression (Figure 6b-e). We present these genes, scored by several unbiased approaches with the introduction of the biology-based metric, as the prioritized biomarkers and subject them to further validation using available human datasets.

### Prioritized biomarkers predict disease severity in validation on human RNA-seq datasets

We aimed to evaluate the prioritized biomarkers in available human hepatic RNA-seq datasets obtained from liver biopsies (Figure 7). First, differential expression analysis identified significant biomarker candidates in these datasets. Then, naïve Bayes-aided feature selection was applied to find the best-performing gene sets, which were then utilized to predict disease severity with random forest.

**Figure 7.**
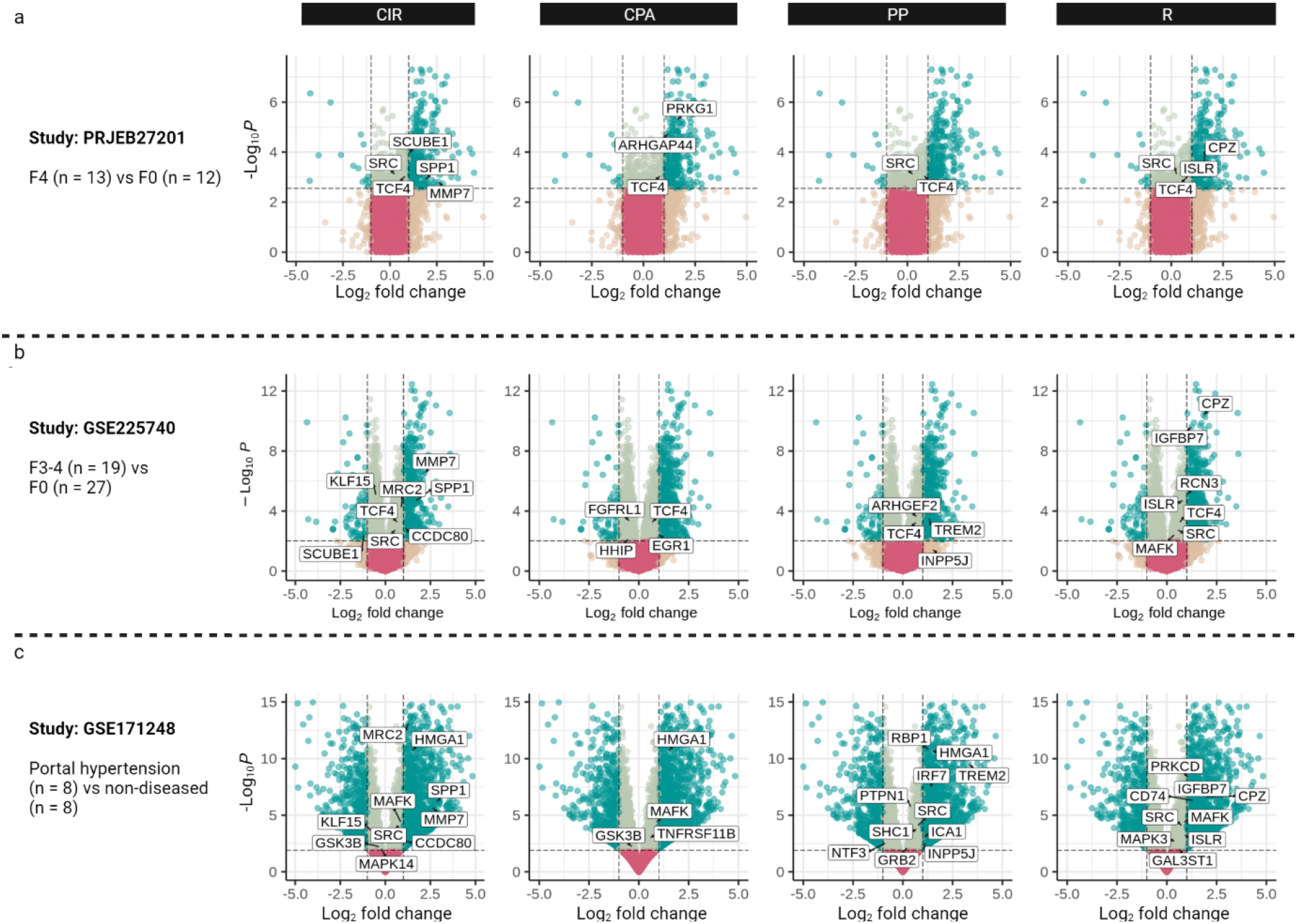
Validation of the prioritized biomarkers in human liver disease datasets. Only genes that are significantly differentially expressed between study groups are present. The comparisons were performed between **(a)** patients with histological cirrhosis (F4) and non-diseased; **(b)** patients with advanced fibrosis/cirrhosis (F3-F4) and non-diseased; **(c)** patients with portal hypertension and non-diseased accordingly. PH = portal hypertension.

In the first analysis, comparing F4 (cirrhosis) and F0 (no fibrosis), the most extensive molecular set was related to our cirrhosis core genes, followed by CPA, regression, and PP core genes (Figure 7a). *SCUBE1*, *PRKG1,* and *CPZ* were the most robust upregulated biomarkers.

The second validation used another dataset of patients with F3-4 and F0 histologic stages. Notably, *SCUBE1* was again the cirrhosis biomarker present there; however, this time, it was downregulated, unlike the first analysis (Figure 7b). In our own dataset, *SCUBE1* showed upregulation in CIR, followed by downregulation in R1+R2, thus, rather supporting an ‘upregulation’ role of SCUBE1 in F4 patients. *CPZ* and *IGFBP7* were the most upregulated regression biomarkers detected from our regression core genes.

The final analysis included patients with cirrhotic portal hypertension and non-diseased controls. Despite the modest number of included patients, PP biomarkers among DE genes were more profound than in previous datasets; *HMGA1*, *TREM2*, *RBP1,* and *IRF7* were the most upregulated biomarkers. There were also regression biomarkers not significantly DE in the earlier comparisons: *CD74* and *MAFK* (Figure 7c).

The machine learning-based feature selection prioritized gene sets, ranging from four to eight genes in size. *CPZ*, the regression biomarker, was included in all best-performing sets. In prediction, the highest score was achieved with *CPZ* (R biomarker), *HMGA1* (TF for cirrhosis, CPA, PP), *IRF7* (TF for PP), and *MRC2* (cirrhosis biomarker), reaching the accuracy of the random forest of 0.933 (Supplementary Table 1).

These findings strongly underline the potential clinical value of the prioritized biomarkers inferred using established animal models. Further validation with appropriate patient cohort composition (e.g., including available PP and CPA readouts, with expected regression groups) is, however, required.

### Cell type deconvolution suggests the contribution of hepatocyte, endothelial, hepatic stellate cell, and macrophage signatures in fibrosis and its regression

Deciphering variations in cell composition across study conditions may give valuable insights into disease pathophysiology. However, due to the complexity of sample preparation and equipment requirements, single-cell methods have yet to become broadly used in the hepatology field. Here, we applied supervised deconvolution to identify the presence of cell-type signatures. Two datasets for signature extraction were selected: single-cell RNA-sequencing (scRNA-seq) of mononuclear macrophages from mouse liver tissue treated for four weeks with CCl_4_ (scCCl_4_)^3^ and single-nuclei RNA sequencing (snRNA-seq) of frozen mouse liver tissue treated with TCDD (snTCDD)^40^ (n = 24) (Supplementary Figure 7-1).

Deconvolution analysis resulted in the macrophage signatures from the sorted scRNA-seq dataset and both parenchymal and non-parenchymal cells from the scCCl_4_ dataset. The two groups of identified Kupffer cell signatures (KC) had various functional markers: (1) cells expressing *Cd63* and *Lgals3bp*, and (2) cells expressing *Vsig4* and *Marco* (Supplementary Figure 7-2a). Among other cell types, HSC and hepatocytes (with periportal markers: *Ass1*, *Pck1*, *Slc7a2*, and *Hal*) had the most prevalent signature (Supplementary Figure 7-2b).

The most robust signatures were found for hepatocytes, macrophages, endothelial cells (source: snTCDD) and HSC (scCCl_4_) (Figure 8a). In CIR, hepatocyte signatures were suppressed as compared to HC and regression, and the macrophage signature showed a reversed trend. Notably, HSC signatures showed upregulation in CIR and R1, which partially restored in R2, likely suggesting the dynamics of a specific HSC subpopulation. The endothelial signature was upregulated in CIR_TAA_.

**Figure 8.**
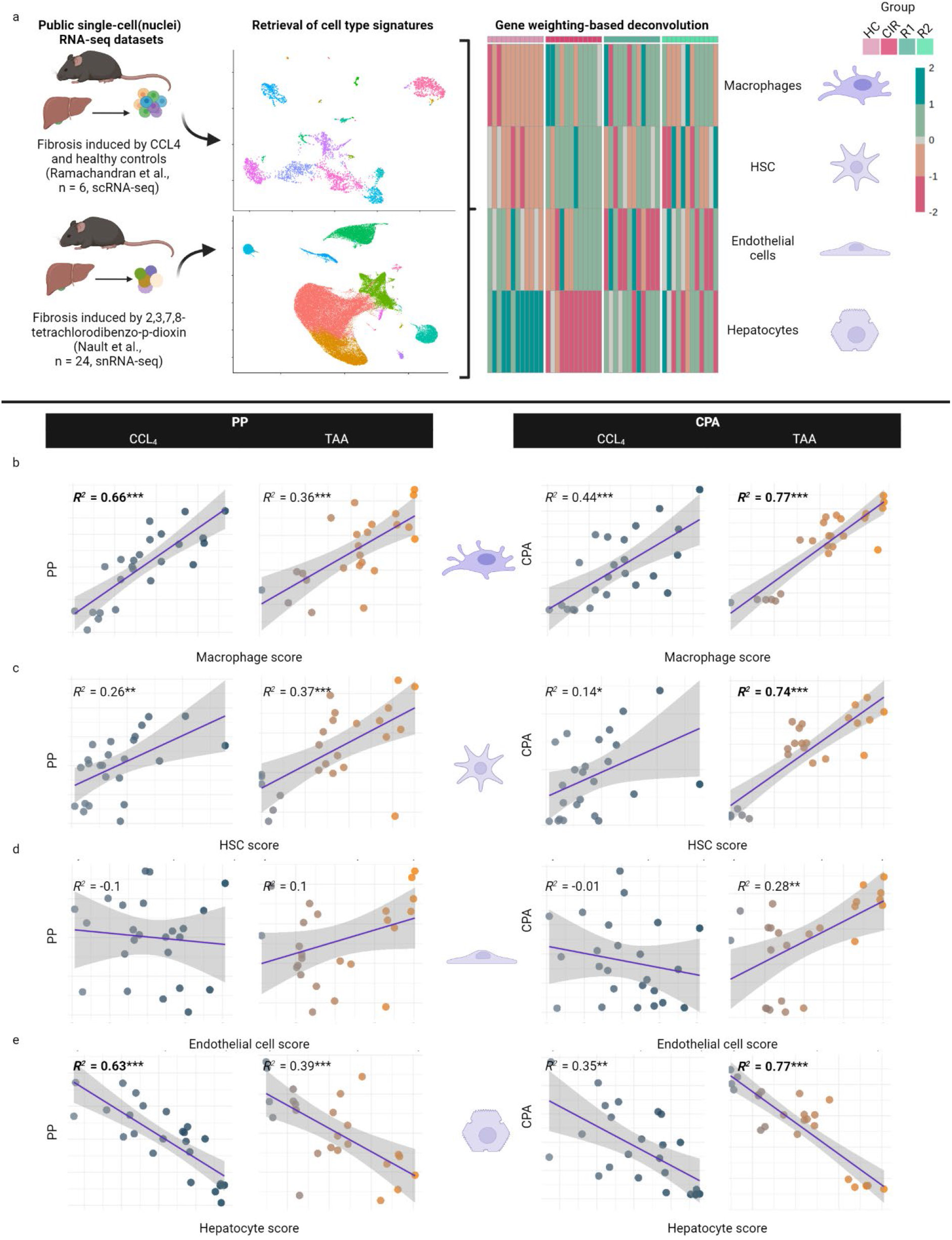
Deconvolution analysis shows cell-specific signatures in mouse RNA-seq datasets and their relationship to PP and CPA. **(a)** Cell type deconvolution of our study dataset using published single-cell datasets. The heatmap illustrates deconvoluted cell types with the most robust signatures. **(b – e)** We found a linear correlation of the respective cell type score to PP and CPA. Cases shown in bold are those in which we consider cell score contributing to parameter variance (R2 > 0.6). The color indicates the respective cell type’s low (grey) or high (blue for CCl4, orange for TAA) score value. HSC = hepatic stellate cells. PP = portal pressure. CPA = collagen proportionate area. p<0.05; ** p<0.01; *** p<0.001.

We performed linear regression to investigate whether there is a linear relationship between these cell scores in each sample with PP and CPA values on the other side. The CPA in the complete TAA dataset positively correlated with all cell scores (Figure 8b-e). However, only macrophage and hepatocyte scores were linked to CPA in CCl_4_. PP significantly correlated with all cell scores in HC_TAA_ and CIR_TAA_, but this relationship did not follow with the inclusion of regression groups.

We further explored the expression of genes with the highest cirrhosis score and biomarker candidates for cirrhosis and CPA on a single-cell resolution in the above mentioned datasets. Cholangiocytes, macrophages including Kupffer cells, and dendritic cells showed the most robust signature with high cirrhosis score (Supplementary Figure 7-3, 7-4). Cirrhosis markers were downregulated in hepatocyte populations, and several CPA markers were upregulated in HSC and portal fibroblasts (Supplementary Figure 7-3).

These findings suggest that the functional state of macrophages, non-parenchymal cells, and hepatocytes are linked to portal hemodynamics and CPA.

## Discussion

In this study, we profiled the hepatic transcriptomes of widely used animal models of liver fibrosis (carbon tetrachloride, CCl_4,_ and thioacetamide, TAA) at key experimental time points of peak fibrosis induction and during one or two weeks of spontaneous disease regression.

Liver disease readouts showed that the CCl_4_ model had more pronounced portal hypertension, liver injury, and fibrosis over the same time course. Yet, both models showed robust transcriptomic features of liver fibrotic such as matrix turnover, with collagens *Col5a2*, *Col1a1*, and Fibrillin 1 being highly upregulated, cell adhesion, and macrophage activation. The common downregulation involved metabolic pathways and PPAR, for which pharmacologic agonists are currently being investigated in clinical trials^28, 41^.

We also described models’ unique signatures and noted differential regulation in delta-Notch, Wnt, and Shh pathways, which could be considered for testing therapeutic strategies targeting their down- or upstream signalling. Despite having less pronounced liver disease readouts, CIR_TAA_ demonstrated transcriptional signs of integrin-regulated cell adhesion, which is notable in the context of the ongoing clinical trials with specific integrin inhibitors involving patients with biliary fibrosis^42^.

Cessation of liver injury for one week already resulted in significant upregulation of previously suppressed pathways in fibrosis, such as cytochrome family proteins, bile acids, and amino acid pathways (Supplementary Figure 3-1). There were, nevertheless, only a few genes differentially expressed between one and two weeks of regression. However, disease remnants’ signature was still present, such as *Col5a3* and *Fbxo31* (indicators of the HSC activation) among upregulated genes in TAA. These findings support that, in future studies, regression periods should be split between early (cessation-related) and delayed regression to better distinguish a rather long-term fibrosis regression signature from tissue repair and metabolic recovery processes.

Solute carriers, cytochrome, and metabolic enzyme genes represented a large group perturbed by the fibrosis induction (e.g., *Cyp4a12a*, *Cyp2j5*, *Slc1a2*). Some of them were among high-ranked potential biomarkers, such genes as *Akr1c18* and Slc9a5 (associated with portal pressure), or *Slc35c1* and *Cyp1a1* (regression). Further mechanistic studies are required to determine whether their pharmacological activation may relieve the metabolic phenotype following experimental fibrosis (suppression of bile acid, amino acid, and eicosanoid turnover).

We revealed that differentially expressed genes could be combined in four dynamic patterns, with either up-regulation or down-regulation in fibrosis and then up- or down-regulation in regression. Notably, the groups following double upregulation or downregulation across the two models were smaller; this might indicate both a long-term effect of fibrogenesis or shifts in tissue homeostasis that would require therapeutic support to recover. A time-series experiment with transcriptomic profiling of liver biopsies in experimental fibrosis should be considered to provide a definitive answer.

Two independent methods (WGCNA and TMLE) revealed genes strongly linked to the liver disease readouts (i.e., portal pressure, collagen proportionate area, ALT, AST) or the fibrosis/regression status. For example, *Nucb2*, not previously studied in the portal hypertension context, was involved in the development of experimental arterial hypertension in obesity^43^; Moreover, we identified that *Cd74* and *Igfbp7* seem to regulate early regression response and persist as upregulated through both weeks. The latter was already proposed as a therapeutic target in NAFLD, yet, its mechanistic role remained elusive^44^. Overlapping results of these two methods prioritized biomarker candidates having both well-described and understudied roles in liver disease.

We also found that, from more than 1300 known transcriptional regulators in mice, 58 were perturbed in our models. Next to the nuclear receptors, we identified as significant those regulators whose role in liver diseases is primarily unknown, such as *Creb3l3, Akap8l, Tcf4,* and *Usf2*. Some TFs were included in the regulation of most of the biomarker candidates (*Hmga1*, *Tcf4*, *Mafk*), thus potentially revealing them as key regulators of transcriptional changes in cirrhotic portal hypertension.

Analysis of the multi-layer network, constructed from the prioritized biomarkers, their regulators, and database-mined interaction targets, led to identifying a few dozen hub genes, which were then further validated in human RNA-sequencing datasets, showing a profound presence in human cirrhotic portal hypertension and achieving promising disease severity prediction in all datasets, with better accuracy in cirrhotic portal hypertension. Despite the availability of the RNA-seq data from human patients stratified according to the fibrosis severity, missing portal pressure and liver injury readouts limited our opportunities for human validation. Hence, the transcriptional biomarker candidates should be validated in a study with such readouts available, prioritizing those we validated in the presented datasets. Profiling of hepatic transcriptomes from patients with cured liver disease, such as after sustained virologic response to antiviral therapy in hepatitis C, will shed light on the clinical value of the regression markers identified in this study.

Finally, cell type deconvolution analysis allowed the identification of signatures of macrophages, non-parenchymal cells, and hepatocytes and their relation to the liver disease readouts. Despite the availability of human scRNA-seq data, employing reference from a similar phenotype to experimental conditions is required to extract accurate cell signatures and thus achieve robust prediction. To further investigate cell contribution and improve accurate cell score prediction, RNA-seq can be paired with FACS for the cell types we could robustly deconvolute (e.g., Kupffer macrophages and endothelial cells) or with low sample-size single-cell RNA-seq for a reference deconvolution.

To summarise, we characterized the hepatic transcriptomes of two commonly used liver fibrosis models, including during a well-defined regression phase, and thereby identified their functional similarities and differences. We identified transcripts with the potential to be developed into prognostic biomarkers or considered for therapeutic applications. Our transcriptomic dataset allowed us to select candidate biomarkers strongly associated with fibrosis progression and regression or key liver disease severity readouts, such as portal pressure or ALT levels. The computational analysis provided insights into their regulatory networks and prioritized genes suitable for screening and assessment in human cohorts. Further mechanistic studies and validation experiments using human liver tissues of well-characterized patient cohorts are warranted to confirm our experimental findings.

## Materials and Methods

### Animal models of liver fibrosis and regression

All experimental animal procedures were approved by the Austrian Federal Ministry of Education, Science and Research (BMBWF-V/3b/2020-0.398.007). Male C57BL/6JRj mice were housed and handled according to the standards of care and maintenance of the Center for Biomedical Research at the Medical University of Vienna. Animals were fed standard autoclaved rodent chow and had water access *ad libitum*. Fibrosis was induced by the administration of carbon tetrachloride (CCl_4_) or thioacetamide (TAA). CCl_4_ was gavaged as a 20% v/v solution in olive oil at a dose of 2 mL/kg body weight three times a week. TAA was injected intraperitoneally in 0.9% saline at a dose of 150 mg/kg thrice a week. The dose was gradually increased over the first week for CCl_4_ administration and over the first two weeks for TAA administration. The total time of fibrosis induction was 12 weeks, followed by one or two weeks of regression. A total of 48 C57BL/6JRj male mice with complete datasets on key liver disease readouts were selected for this study. The experimental cohort was divided into two different fibrosis models (CCl_4_ and TAA) of 24 animals each (Figure 1a). Each model comprised four groups: healthy control with vehicle treatment (hereafter HC), positive control at the peak of the 12-week induction by CCl_4_ or TAA respectively (CIR), one week of regression post-induction (R1), and two weeks of regression (R2) upon cessation of the stimulus (Figure 1b). Each group included six animals.

### Key liver disease readouts

The following biological parameters were assessed in each animal: portal pressure (PP), collagen proportionate area (CPA), serum levels of alanine aminotransferase (ALT) and aspartate aminotransferase (AST) (Table 1). The fibrosis assessment was performed by digital semiautomated histomorphometry of the relative fibrotic area on full liver lobe scans as previously described^45^. Briefly, the left lateral lobe of the liver was harvested, formalin-fixed, and paraffin-embedded. The 4 µm sections were stained by picrosirius red with fast green counterstain and scanned with Olympus VS200. The whole-slide images were then analysed using HALO (V3.3.2541.184, Indica Labs), and the percentage of collagen-positive staining (% CPA) was quantified with the Area Quantification module. PP was measured by direct cannulation of the portal vein under anaesthesia (ketamine 100mg/kg, xylazine 7.5mg/kg), using our established hemodynamic setup^46^. Serum levels of ALT and AST were measured as established surrogates of liver injury.

### Transcriptomic assessment with RNA-sequencing

Left and right median liver lobes were harvested and immediately snap-frozen. RNA extraction from the right median lobe was performed using the RNeasy Mini Kit (Qiagen, Germany). RNA concentration and purity were accessed with NanoDrop One (Thermo Fisher Scientific, MA, United States) and submitted to the Biomedical Sequencing Facility of the CeMM Research Center for Molecular Medicine of the Austrian Academy of Sciences for further processing. The quality of the RNA was assessed with 2100 Bioanalyzer (Agilent, CA, USA), and RNA integrity ≥ 7.0 was used as the threshold (median integrity number: 7.65). The library preparation was done with a TruSeq Stranded mRNA LT sample preparation kit (Illumina, CA, USA). Samples were diluted, pooled into next-generation sequencing (NGS) libraries in equimolar amounts, and sequenced on Illumina HiSeq 4000 Systems (single-end, 50bp). The pipeline for base calling, QC, and demultiplexing was based on Picard tools (version 2.19.2) (Supplementary Figure 1). The alignment was performed to Mus musculus GRCm38/mm10 Reference Genome Assembly^47^. The quality controls included sequencing depth evaluation, mapping percentage, and outlier detection with RNAseqQC (v. 0.1.4). Genes with no variance between samples, and those detected in fewer than 10% of the samples, were removed from the analysis. Gene annotation was performed with AnnotationDbi (v. 1.60). Dimensionality reduction was performed with PCAtools (2.10). RNA-sequencing produced an average of 27 487 574 transcript reads per animal and a reference mapping rate of 98.2%. The annotation of reads resulted in 20 869 identified genes that were utilized in the subsequent analysis.

### Differential expression (DE) analysis

DE was performed using the DESeq2 package with a detection limit of 2 counts^48^ (v. 1.38.1). Count normalization with variance stabilizing transformation was performed. The Wald test was conducted with a design formula of ∼ Model + Group for the pairwise analysis with ashr shrinkage (v. 2.2-54). The absolute log2FC > 1.5 was used as a cutoff to determine the direction of differential expression. The multiple-group comparison was performed with the likelihood ratio test (LRT) in the reduced (intercept-only; ∼ 1) model. An adjusted p-value cutoff of < 0.01 was applied for both types of analysis to determine significant genes.

The Wald test was applied to characterize the models, compare groups, and validate the findings. The LRT test output provided a list of all genes significantly perturbed among all study groups, and this was utilized in pseudotemporal analysis to define the perturbation scores (described in own section).

For targeted analysis of genes involved in matrix turnover among identified DE, genes included in the Reactome pathway R-HSA-1474244 were utilized.

### Gene set analysis

We assessed gene sets using fgsea^49^ (v. 1.24.0) and enrichR^50^ (v. 3.2). For fgsea, genes were ranked based on the formula: 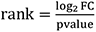 and gene sets of WikiPathways^51^ (2019, mouse) were used. In gene lists where such ranking was not possible (e.g., biomarker modules, network nodes), we used a hypergeometric test of EnrichR^52^ with the MSigDB^53^ hallmark and Reactome pathways, indicating the database in figures legend. Only pathways with a q-value ≤ 0.25 are reported. Usage of another threshold, or subsetting the top-ranked pathways, is noted in figure legends.

### Pseudotemporal dynamics and perturbation scores

DE genes from the LRT test were used for pseudotemporal analysis. Study groups were defined as pseudo time points, and 3000 genes with the highest significance were clustered using DEGreport (v. 1.35.0). This allowed the identification of groups of genes with similar dynamics in the direction of fibrosis and regression.

Perturbation scores were calculated to determine the impact of cirrhosis and regression on gene expression. The cirrhosis score was defined as the impact of cirrhosis with HC as a baseline, and the regression score accounted for CIR as a baseline towards regression. First, stepwise regression fit was performed with maSigPro^54^ (v. 1.70.0) for genes differentially expressed (DE) with the LRT test, resulting in goodness to fit (R^2^) for each gene. Benjamini-Hochberg correction was used with a threshold of padjusted < 0.01. Next, perturbation scores were calculated based on the formula:

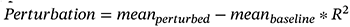

where meanperturbed denotes the mean expression level in cirrhosis or R1+R2. Pearson correlation was used to evaluate the relation between the cirrhosis and regression scores, and pathway analysis was performed. These scores were used for later hub gene identification.

### Biomarkers discovery with TMLE and WGCNA

To identify potential biomarkers associated with liver disease readouts, cirrhosis, and regression, we employed two statistical approaches. First, CCl4 and TAA datasets were merged and vst-normalized for input for WGCNA^55^ (v. 1.71) for gene-readout correlation analysis. The network was constructed using a signed type with a defined minimal module size of 10 genes and an optimal soft threshold of 5. All modules with a trait correlation strength > 0.6 and p < 0.01 were considered for further evaluation. We ranked genes separately for each readout (gene significance score). Finally, the upper and lower 10% genes were subsetted by this rank, given that both the strongest positive and negative gene significance indicate a strong association with a trait.

Second, we used the biotmle^56^ (1.22) toolkit as a more conserved approach^57^ to rank genes based on their association with liver disease readouts and study groups, accounting for other covariates. A cutoff of an adjusted p-value < 0.01 was applied, and genes in the upper 10% by the TMLE score were selected.

The candidate genes from both methods were overlapped to identify biomarkers robust to different sources of bias and confounding, resulting in four sets (for PP, CPA, CIR, and R1+R2). The candidate genes in these overlaps were considered potential biomarkers and were used in future analysis.

### Identification of transcriptional regulators and their scoring

To identify transcriptional regulators and their scoring, the CeTF package^58^ (v. 1.9.0) was used with the Reverter algorithm^59^, and an adjusted p-value < 0.01 was the threshold. This approach constructs a co-expression network between DE genes and transcriptional factors (TFs) in two conditions of interest (e.g., HC and CIR) and compares the co-expression correlation in these two conditions. TFs with the most differential co-expression are prioritized, allowing the identification of relevant TFs based on their target genes, even if these TFs are not differentially expressed themselves. The curated list of mouse transcription TFs^60^ and DE genes identified in the previous analysis with the Wald test were used for that.

We used GENIE3^61^ to narrow down the identified TFs further, focusing only on those relevant to the animals’ phenotype by scoring the TFs’ interaction with biomarker overlaps for PP, CPA, CIR, and R1+R2. The Extra-Trees algorithm was preferred to the random forest to avoid subsampling bias and to include all possible TFs-genes combinations. The background interactions were defined as TFs-genes interactions with GENIE3 weight < 0.05 and removed.

### Identification of hub genes via multi-layer network analysis

We constructed a multi-layer network for hub gene identification, consisting of the following layers (bottom-top):

- Gene regulatory network (GRN) – prioritized TFs served as network nodes;
- Gene co-expression network (GCN) – biomarkers from overlaps as nodes;
- Protein-protein interaction network (PPIN) – inferred targets of the GCN layer as nodes.

The GRN-GCN inter-layer edges were based on TFs-genes interactions identified with GENIE3 analysis. The GCN layer had intra-layer edges, to obtain which we performed co-expression analysis using WGCNA’s similarity over topological overlaps with signed type and power = 5.

To construct the PPIN, we first used OmnipathR^62^ (v. 3.2.0), and for each gene of the GCN the interaction target was identified using platform’s *omnipath*, *kinaseextra*, *pathwayextra*, *ligrecextra*, and enzyme-substrate databases. These target proteins served as PPIN nodes, and interactions were added as GCN-PPIN edges. Secondly, we added intra-layer PPIN interactions from STRINGdb^63^ (v. 2.10.0), with the experimental score threshold ≥ 0.7 without the addition of new proteins. When the experimental score was unavailable, a STRINGdb confidence score of ≥ 0.7 was applied.

The resulting multi-layer network was imported into Cytoscape (v. 3.9.1) for visualization and scoring with CytoHubba^64^.

We then identified four network cores (NC), for each biomarker sets: PP, CPA, CIR, and R1+R2. For each of them, the GCN subset with respective biomarkers was used together with all GRN and PPIN nodes connected to them, including intra-layer connections of PPIN. Genes for each NC were prioritized via scoring based on a combination of biological knowledge, the double screening method^65^, and machine learning.

For biological knowledge, we used previously calculated perturbation scores (cirrhosis and regression scores) and gene significance for the PP and CPA.

In the double screening method, the maximum neighbourhood component (MNC) and density of maximum neighbourhood component (DMNC) scores were selected. The reasoning behind that was that MNC is a global network score that measures the size of the largest connected component that a node belongs to, while DMNC is a local network score that measures the density of the subnetwork consisting of a node and its neighbours. We used either one of them (as described in the formulas), or two where applicable. By using MNC as a pre-filtering step, the most essential and highly connected nodes in the network were prioritized. DMNC was used to further refine the selection of the genes most relevant to the nodes mentioned above.

Random walk with restart (RandomWalkRestartMH, v. 1.18.0) was prioritized from machine learning methods. It initiated on seed nodes, represented as TFs (GRN layer); then, it iteratively transposed to their neighbours (in the GCN layer), until, eventually, reaching the last node (in the GCN or PPIN layers), or resetting and starting a new walk, hence ranking the nodes with the walk probability. We set the probability of restarting to 0.7, τ (the likelihood of reset in a layer other than GRN) to 0, and used the above-mentioned inter- and intra-layer connections as weights.

Consequently, we identified NC with the following formula:

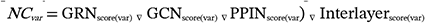

where *del* operator indicates joining operation with preserving inter-layer edges, and *var* denotes one of four biomarker sets. The scores were calculated as following:

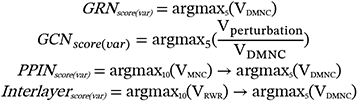

where *V_x_* = set of vertices and their edges for this *var* obtained using one of the indicated scoring approaches; *DMNC* = density of maximum neighbourhood component; *MNC* = maximum neighbourhood component; *RWR* = random walk with restart; *argmax_n_* = function retrieving n components of the list based on the indicated scoring; arrow = chain operation.

If several genes from the argmaxn set had identical scores, all of them were included in the result. If less than five TFs were involved in GRN layer for a biomarker set, all of them were included in the result.

### Validation in human datasets

For validation, human RNA-seq data from the following studies were utilized:

- PRJEB27201: liver biopsies from patients with F4 and F0 (n = 25)^66^.
- GSE225740: liver biopsies from patients with steatohepatitis and non-diseased (n = 74); patients with F3-4 and F0 (n = 46)^67^.
- GSE171248: liver biopsies from patients with cirrhotic portal hypertension and non-diseased (n = 16)^68^

In all scenarios, we investigated the presence of the identified hub genes among DE genes. The Wald test was applied for differential expression with an adjusted p-value threshold of < 0.01. Using Caret^69^ (v. 6.0-94), machine learning algorithms were employed to validate the hub genes, identifying those with the potential for predicting disease severity. Namely, naïve Bayes was preferred for feature selection, considering the involvement of various transcriptional factors in their expression and the assumption that both their regulation and targets follow conditional independence. The candidate biomarker combinations were scored for the classification task (prediction of the above-mentioned disease severity in each dataset), with five repeats of 10 cross-validations applied. Given the human datasets’ modest size and overfitting risks, the random forest in Leave-One-Out cross-validation (mtry = 1) was selected. For each classification task, results included the best set of genes, the accuracy of the random forest-based model, and the Kappa statistic.

### Cell type deconvolution

We utilized MuSiC^70^ and NNLS^71^ algorithms to estimate the cell type proportions in the study dataset for cell type deconvolution. The reference objects were constructed from single-cell RNA-seq (scRNAseq) of CD45-positive cells in healthy and 4-week CCl4 mice models ^3^ (n = 2) and whole liver single-nuclei RNA-seq (snRNA-seq) of models induced with 2,3,7,8-Tetrachlorodibenzo-p-dioxin^72^ (TCDD, n = 24). Only significant DE markers of each cell cluster, identified with Seurat’s FindAllMarkers (parameters: absolute logFC threshold = 1.5, test = Wilcoxon, min.pct = 0.25), were used during deconvolution. MuSiC or NNLS were prioritized depending on their ability to detect cell signatures as non-zero values in all samples. The deconvoluted cell type proportions were compared between groups and models using the Wilcoxon rank-sum test. We used linear regression models with the formula *parameter ∼ predicted cell score* to investigate the association between continuous readouts (PP and CPA) and the predicted scores.

## Acknowledgements

The research group of TR (OP, PK, KB, BSH, ALL, AL, KB, BS, PS) and the study were supported by the Austrian Federal Ministry for Digital and Economic Affairs, the National Foundation for Research, Technology and Development, the Christian Doppler Research Association, and Boehringer Ingelheim. AFR is supported by Angelini Ventures S.p.A. Rome, Italy. The help of Alexander L Lein and Ariton Ljoki with the in vivo experiments is thankfully acknowledged. The authors appreciate the technical assistance of Kerstin Zinober and Martha Seif in this study. The Biomedical Sequencing Facility staff is acknowledged for generating NGS data and related support. Separate figure elements were created with BioRender.com.

## Author contributions

TR, PS, OP, PK, SGK, LP and MT developed the study design. PK, OP, KB, BSH, KB and BS contributed to the animal experiments and readouts measurement. OP performed computational experiments. LPMHdR, AFR and ES mentored the computational part and contributed to the design of additional experiments. All authors contributed intellectual content and participated in creating the final draft of the manuscript.

---

**Supplementary Figure 1.**
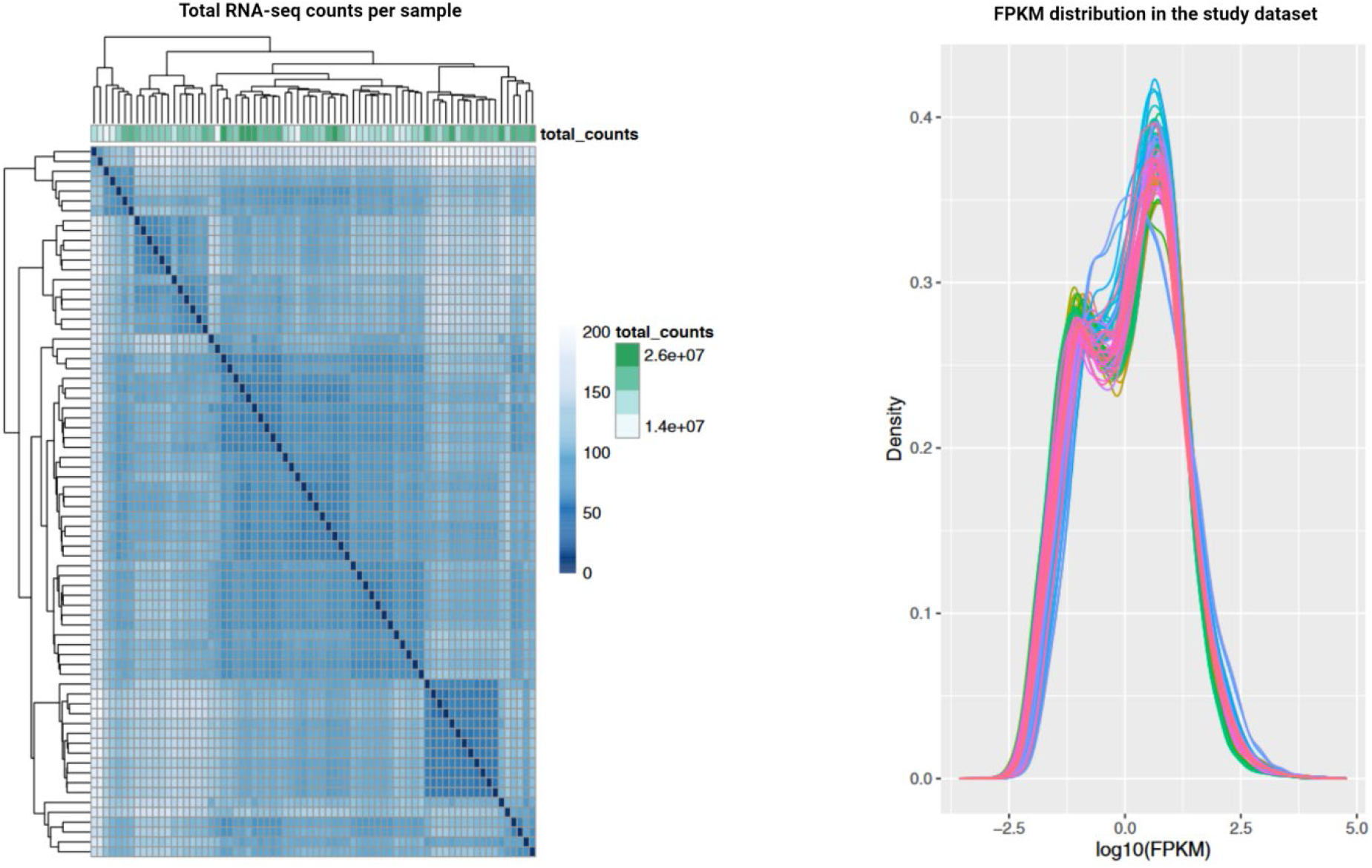
Quality assessment of the generated RNA-seq samples. The heatmap represents the count density per each sequenced sample (represented as rows; n = 48). The density curves represent a distribution of fragments per kilobase of transcript per million mapped reads (FPKM) in each sample.

**Supplementary Figure 2_1.**
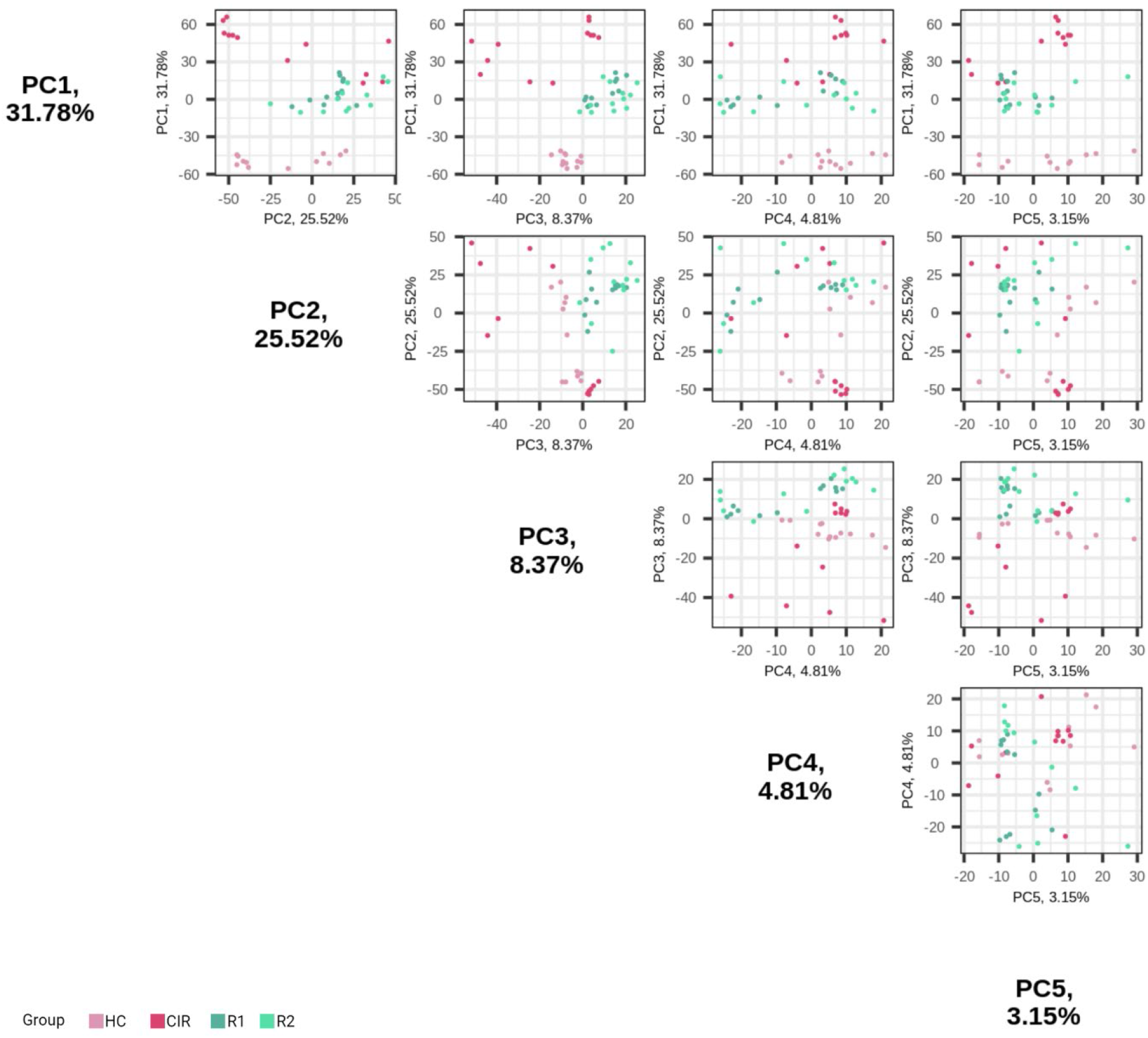
Five most important principal components to distinguish animal groups. The colours are assigned according to the study groups.

**Supplementary Figure 2_2.**
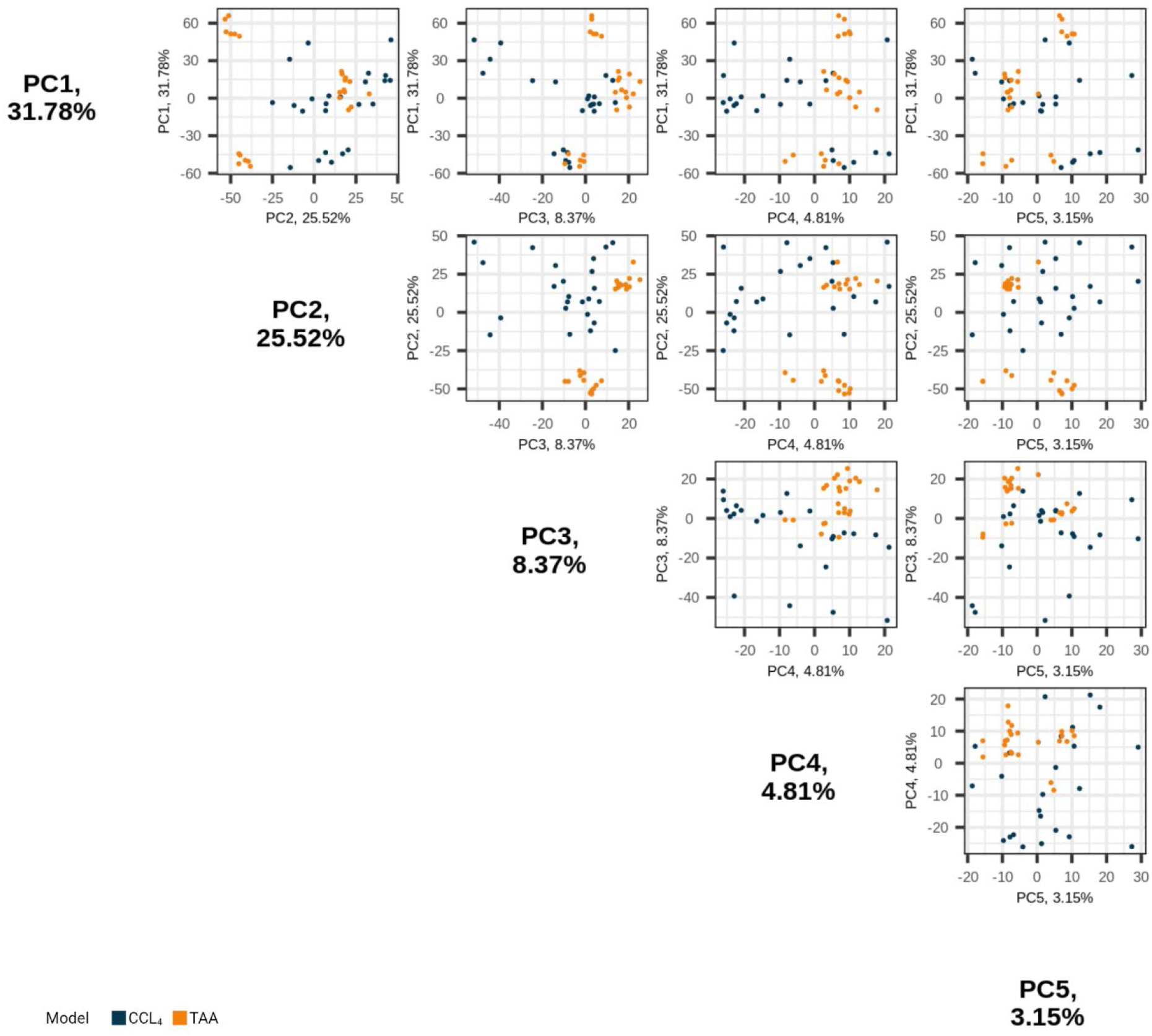
Five most important principal components to distinguish animal models. The colours are assigned according to the animal models.

**Supplementary Figure 2_3.**
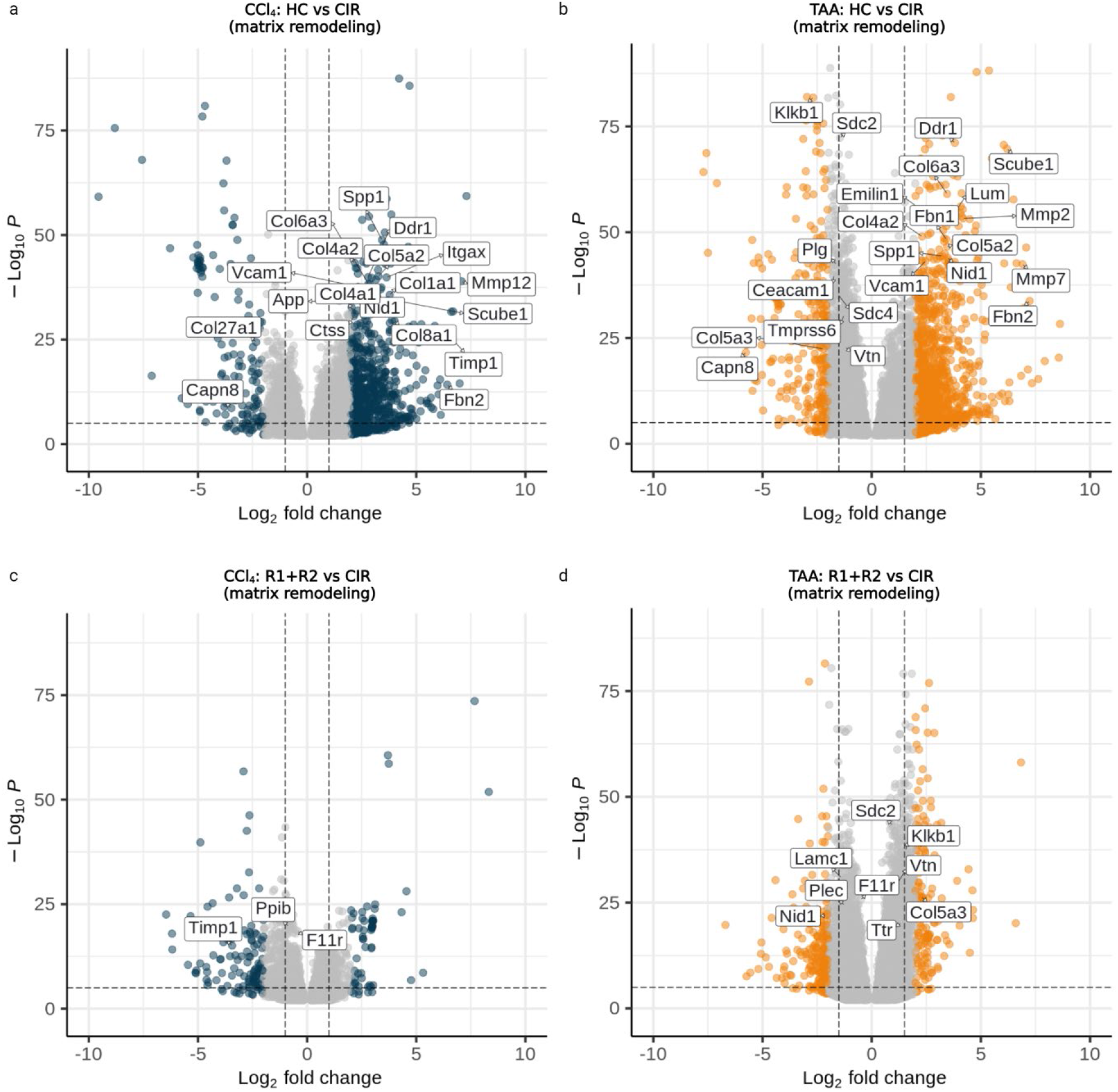
Presence of matrix remodelling genes in differentially expressed genes of the study groups. Comparisons: **(a)** HC_CCl4_ vs CIR_CCl4_. **(b)** HC_TAA_ vs CIR_TAA_. **(c)** R1+R2_CCl4_ vs CIR_CCl4_. **(d)** R1+R2_TAA_ vs. CIR_TAA_.

**Supplementary Figure 3_1.**
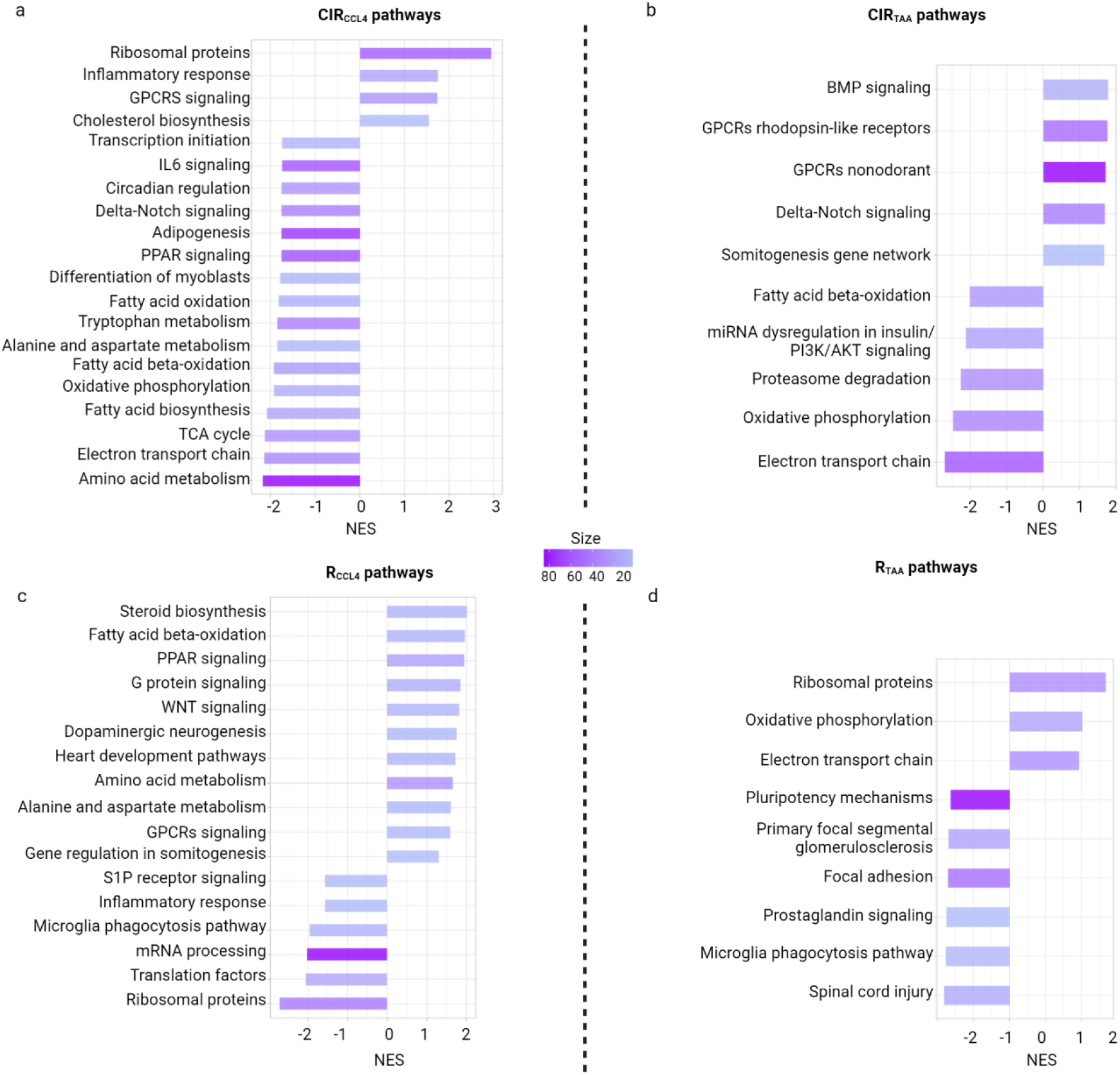
Functional annotation of cirrhosis and regression in CCl_4_ and TAA models. Significant pathways in **(a)** CCl_4_ and **(b)** TAA cirrhosis and regression **(c, d)**.

**Supplementary Figure 3_2.**
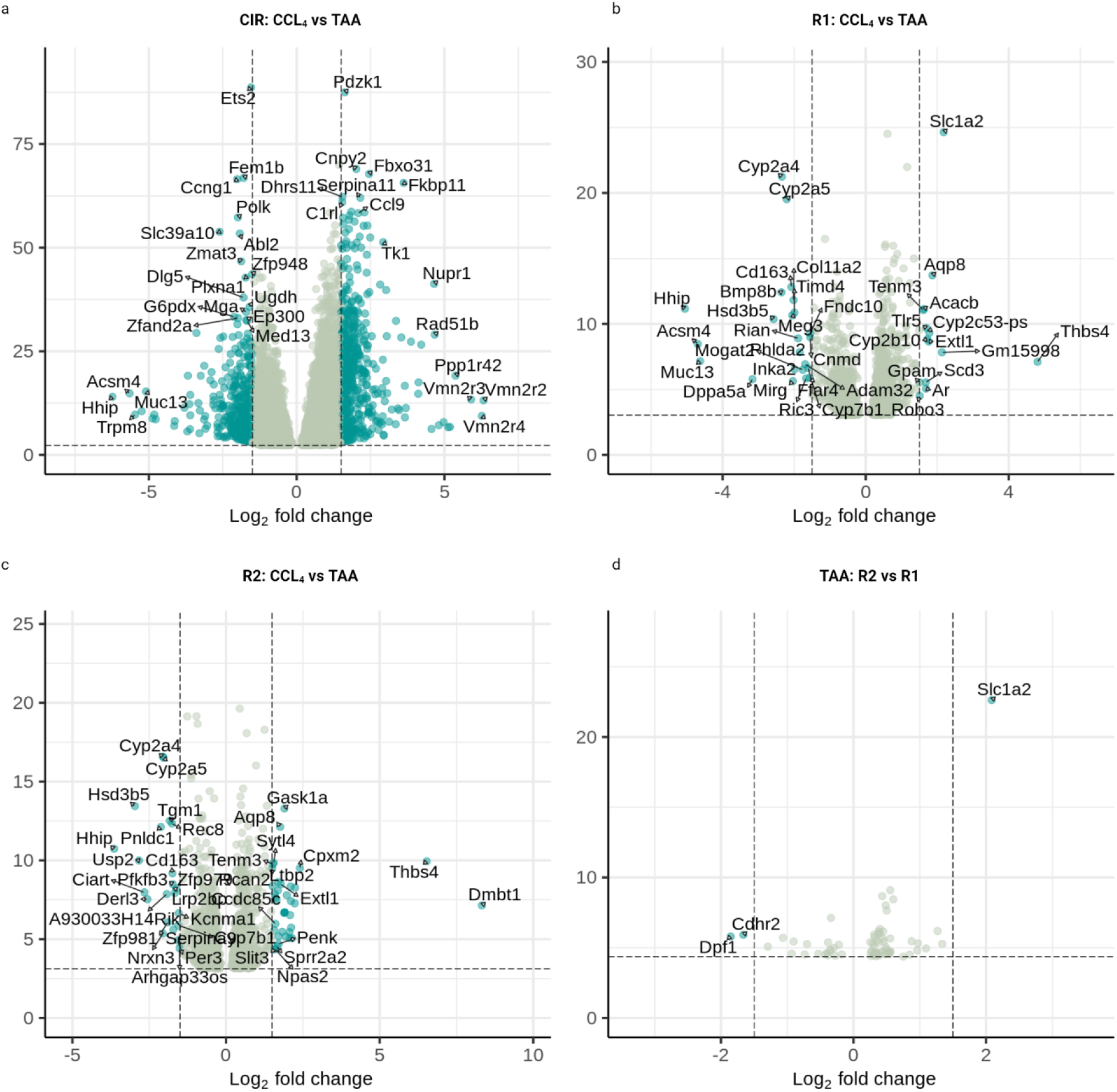
Model-specific differentially expressed markers. **(a)** comparison of CCl4 and TAA features in cirrhosis. **(b, c)** Models feature in R1 and R2 accordingly. **(d)** Differential testing identifies genes significant between R2 and R1 groups in TAA, while no such genes were detected for the CCl4 model.

**Supplementary Figure 3-3.**
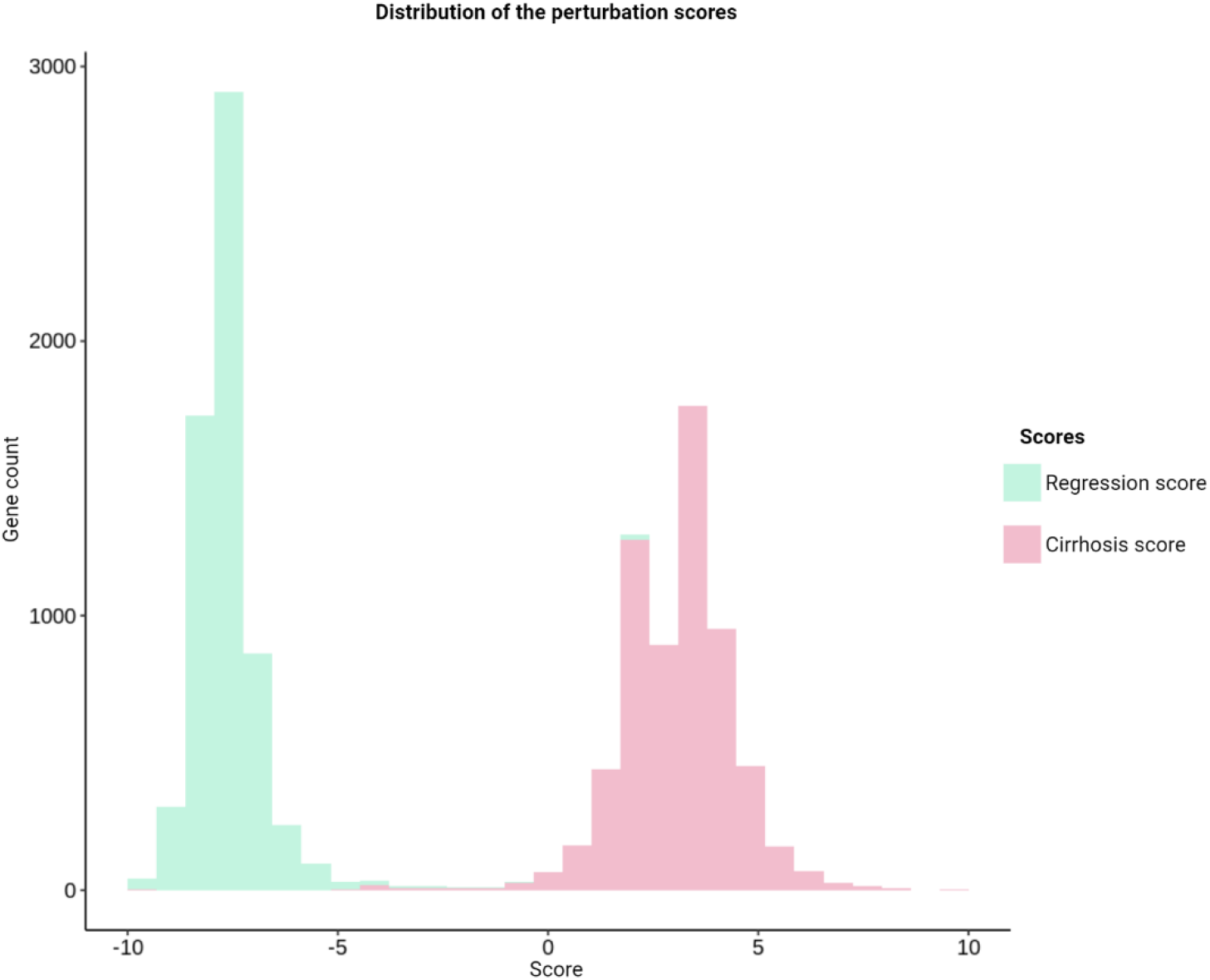
Distribution of the regression and cirrhosis scores. These scores were calculated based on the expression change in this group (CIR or R1+R2) compared to the corresponding baseline (healthy control or CIR) and the coefficient of their linear regression fit to predict CIR or R1+R2 as compared to background genes. Two peaks demonstrate opposite directions in terms of up/downregulation.

**Supplementary Figure 3_4.**
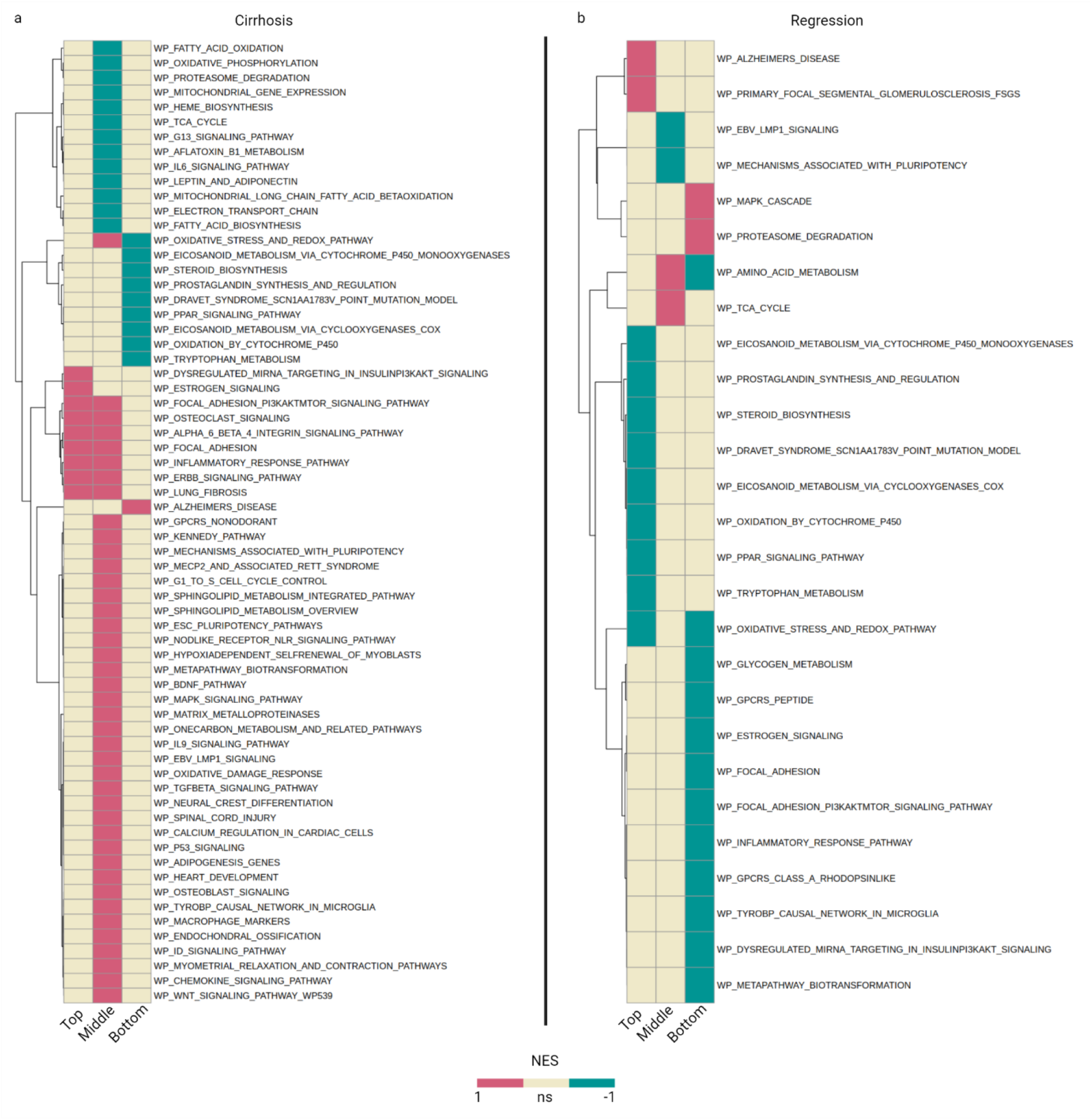
Gene set analysis of gene groups with highest, lowest, and intermediate gene scores. All genes were scored for cirrhosis **(a)** and regression **(b)** gene scores. Each set was then divided into the top (20%), and bottom (20%) scored genes; the remaining genes were marked as “middle”. The colours correspond to the NES of a pathway. NES = normalized enrichment score.

**Supplementary Figure 4_1.**
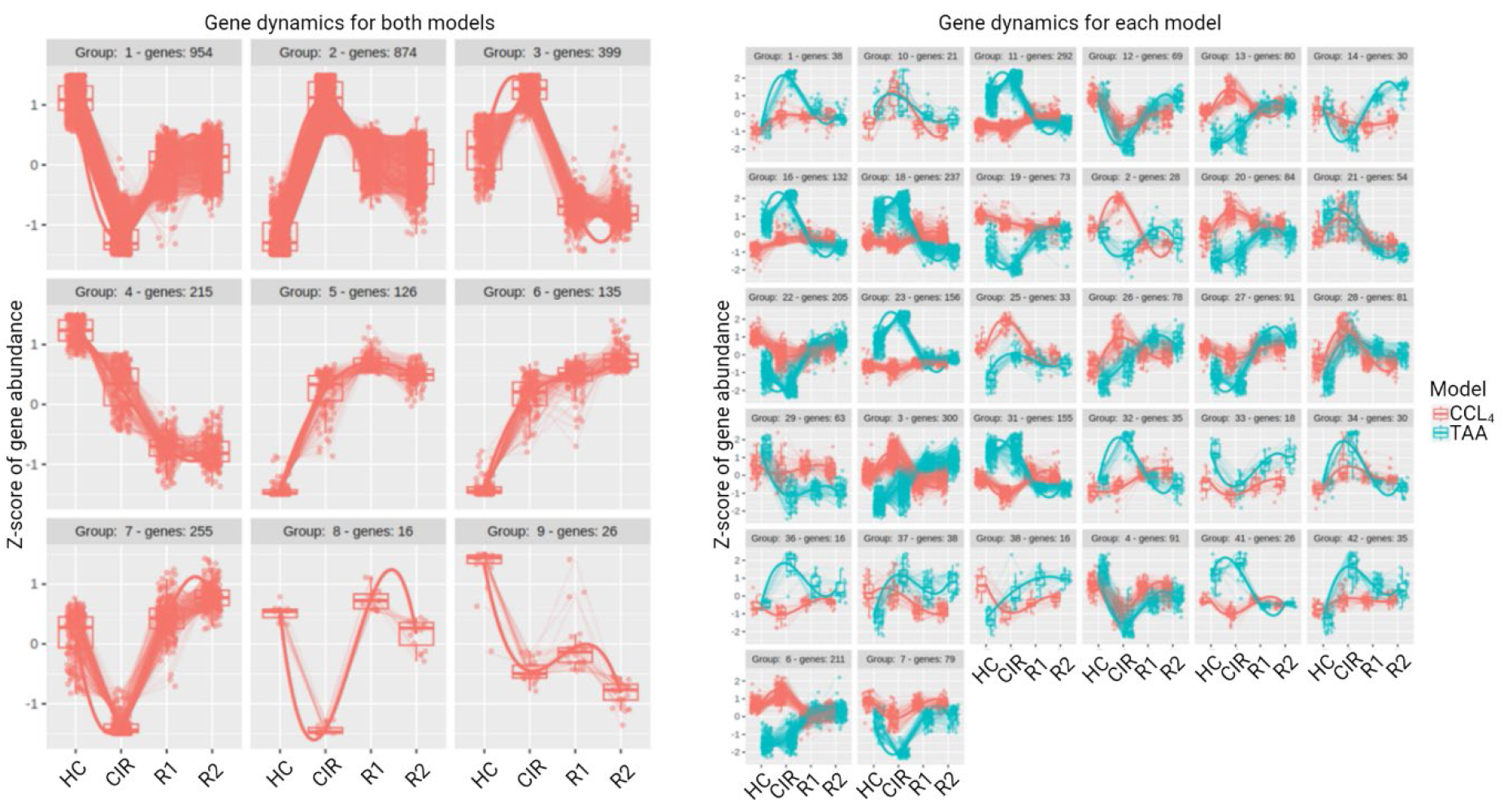
Gene clustering in pseudotemporal points. **(A)** Clusters of both animal batches. **(B)** Clusters of differentially expressed genes in each of the models.

**Supplementary Figure 4_2.**
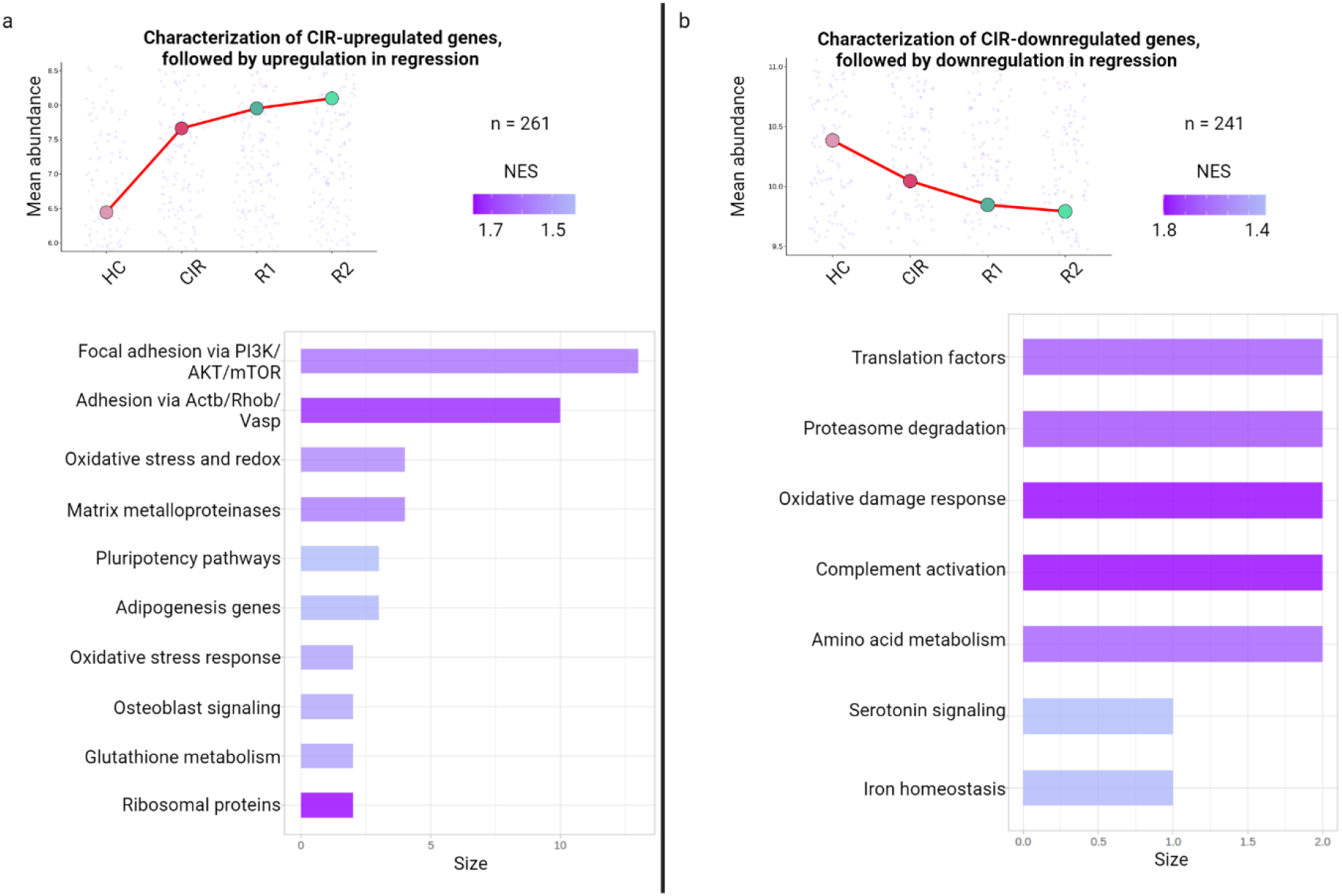
Functional annotation of genes following single-directed dynamics. **(a)** Cluster 3: pathway analysis of genes up-regulated in cirrhosis and upregulated in regression; **(b)** Cluster 4: pathway analysis of genes downregulated in fibrosis and downregulated in regression. No pathways were retrieved with the default adjusted p-value cutoff for Cluster 4, and threshold Padj < 0.35 was used for visualization purposes.

**Supplementary Figure 4_3.**
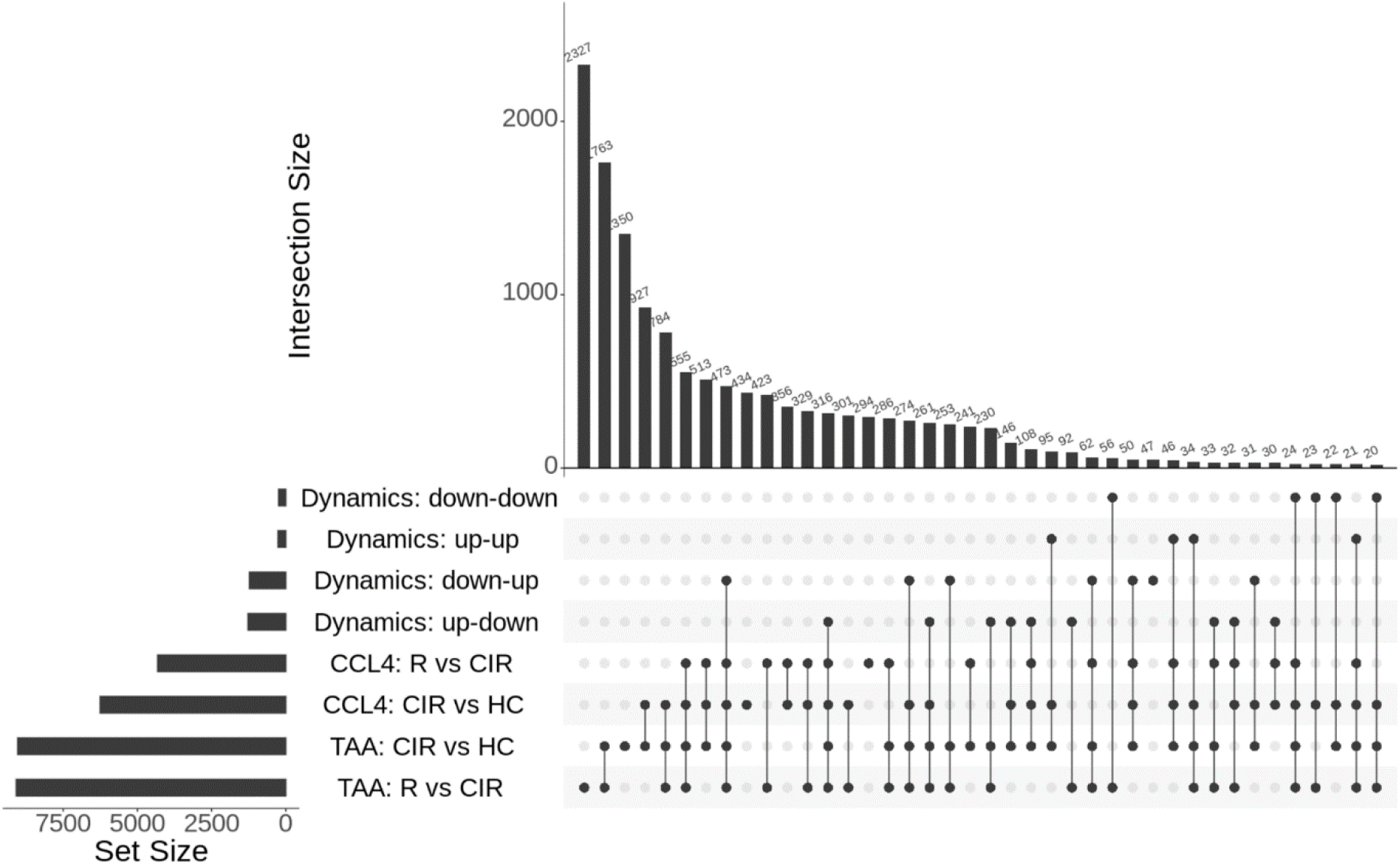
Intersections between groups of differentially expressed genes and gene sets grouped according to their dynamics. The upper bars display the intersection size.

**Supplementary Figure 5_1.**
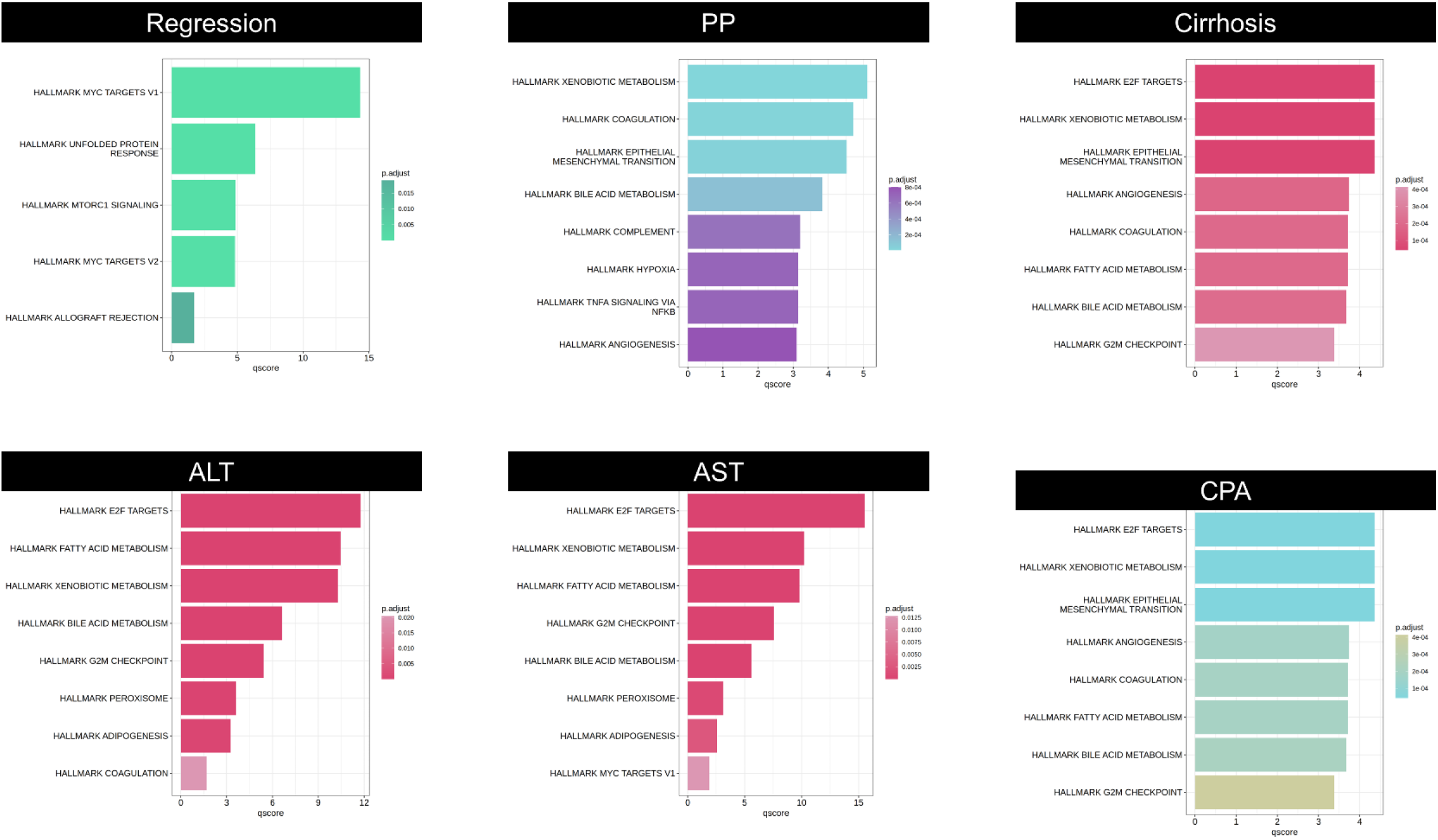
Gene set analysis of genes correlating with key biological readouts. Genes from respective modules were functionally enriched via a hypergeometric test for pathways from the MSigDB Hallmark database.

**Supplementary Figure 5_2.**
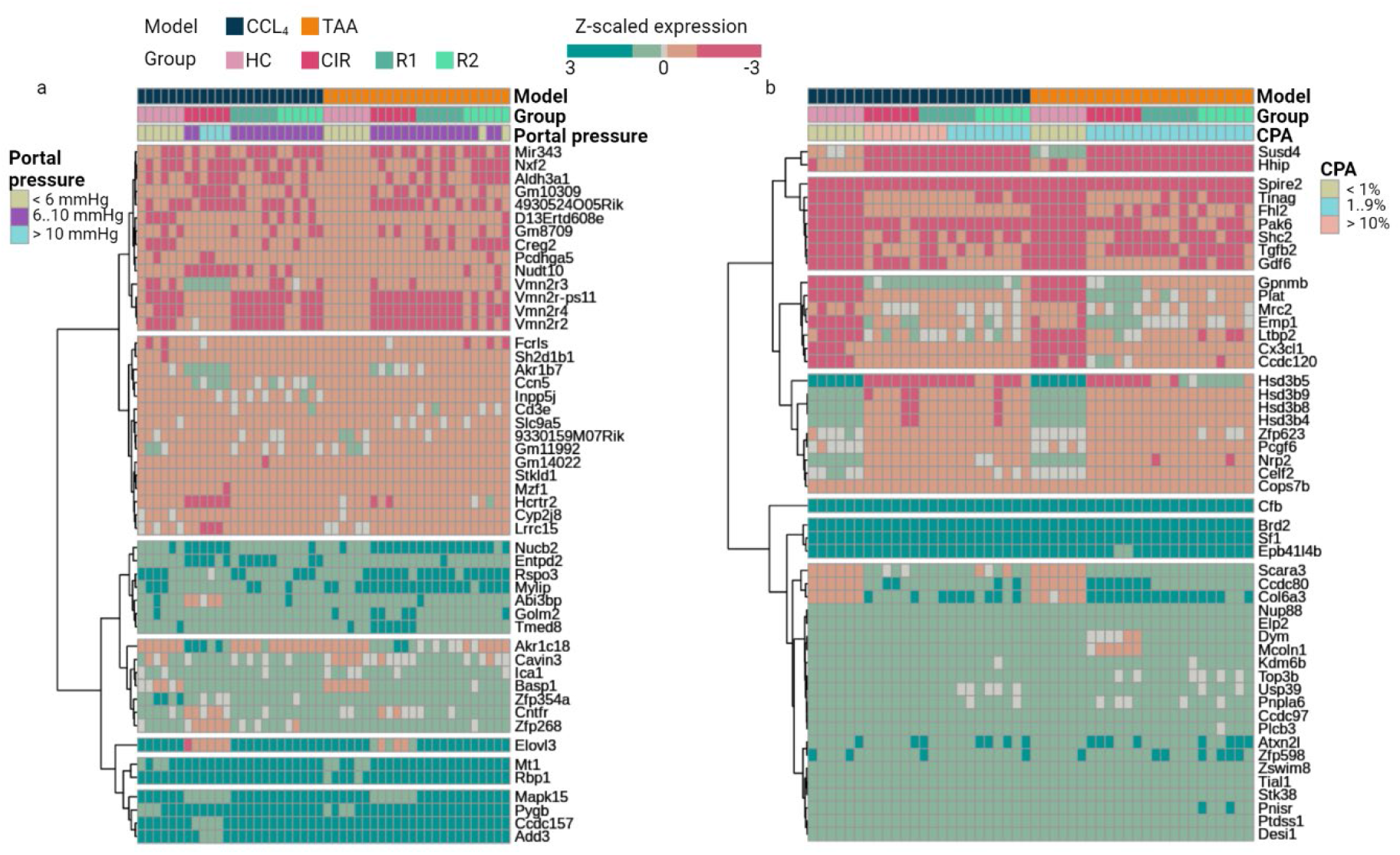
TMLE-prioritized gene associations with biological readouts. Targeted Minimum Loss-Based Estimation genes associated with portal pressure (a) and (b) collagen proportionate area. Fifty genes with the highest adjusted P-value are displayed. Colours indicate expression change.

**Supplementary Figure 5_3.**
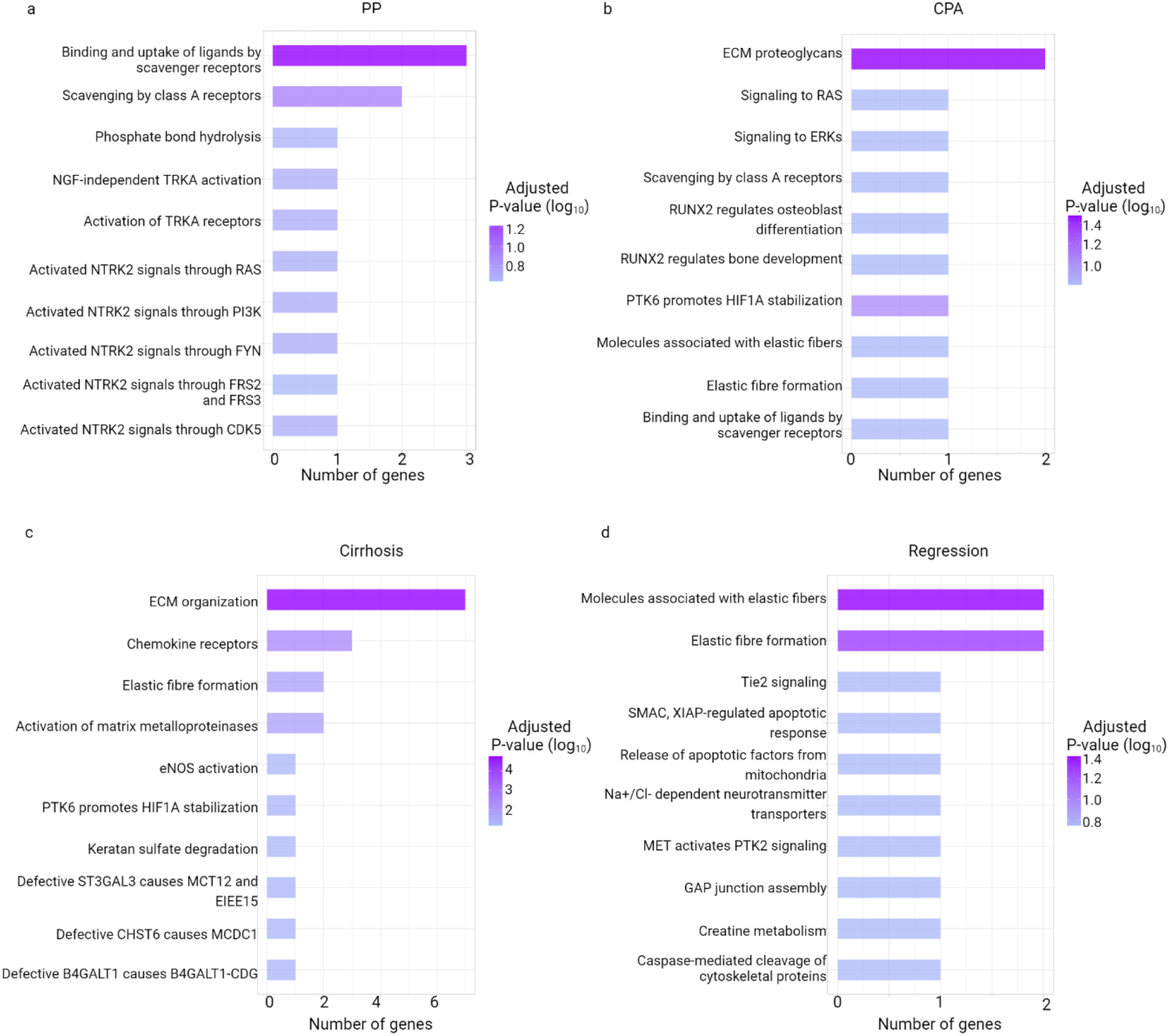
Gene set analysis of the overlaps containing candidate biomarkers. The gene sets are grouped for (a) PP, (b) CPA, (c) cirrhosis, and (d) regression. Enrichment was performed using a hypergeometric test with the Reactome terms.

**Supplementary Figure 6.**
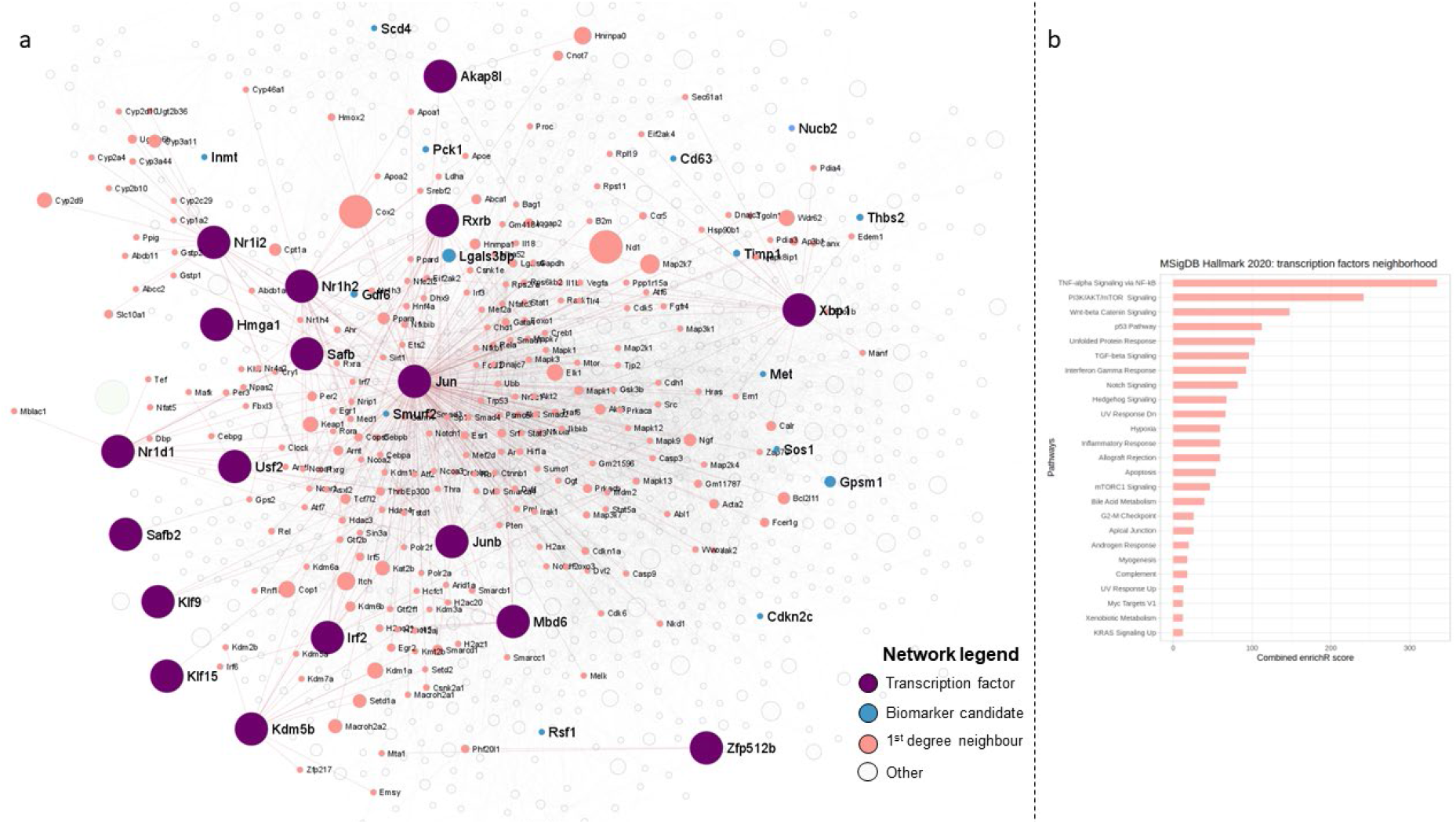
Network analysis reveals signalling niches of transcriptional regulators in fibrosis progression and regression. (a) Identified transcriptional factors (TFs) and differentially expressed genes (LRT test) connected via interaction networks. The first-degree neighbours are highlighted in salmon colour and are of interest to explain the functional importance of the TFs. The blue indicates genes in this network previously discussed in our study. (b) Gene set analysis of the 1st-degree neighbours. Most pathways indicate that identified TFs contribute to inflammation and fibrogenesis. Only significant pathways, identified with MSigDb analysis are presented (adjusted p ≤ 0.05). Vertex size (except TFs) inversely indicates adjusted p-value.

**Supplementary Figure 7_1.**
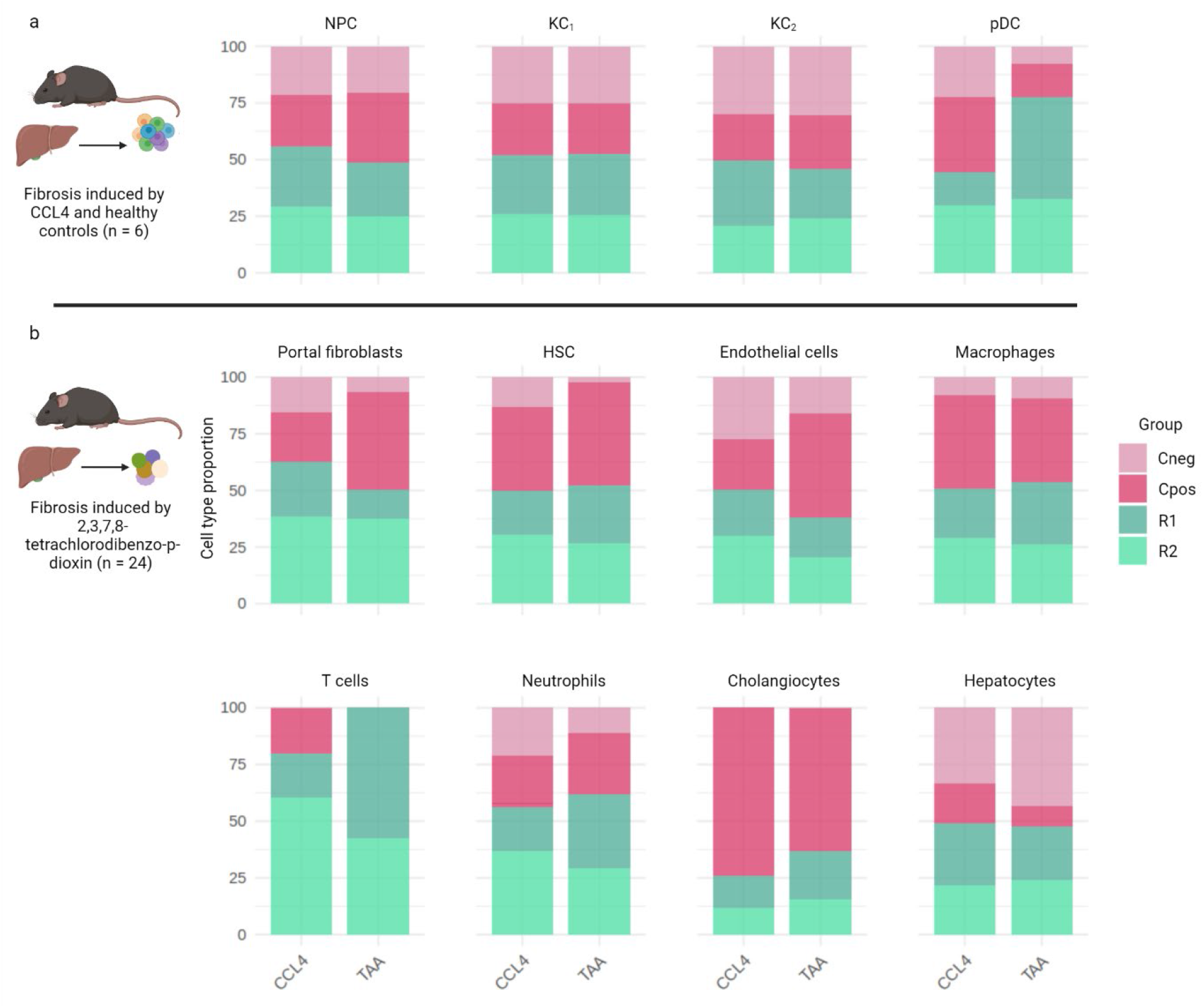
Cell-type deconvolution of bulk RNA-seq data. **(a)** Cell types obtained from single-cell reference (Ramachandran et al., Nature 2019). In the original dataset, the NPC fraction relates to cells expressing HSC and endothelial markers that persisted through CD45+ cell sorting. **(b)** Cell types obtained from single-nuclei reference (Nault et al., Toxicological Sciences, 2022). NPC = non-parenchymal cells. KC = Kupffer cells. pDC = plasmacytoid dendritic cells. HSC = hepatic stellate cells.

**Supplementary Figure 7_2.**
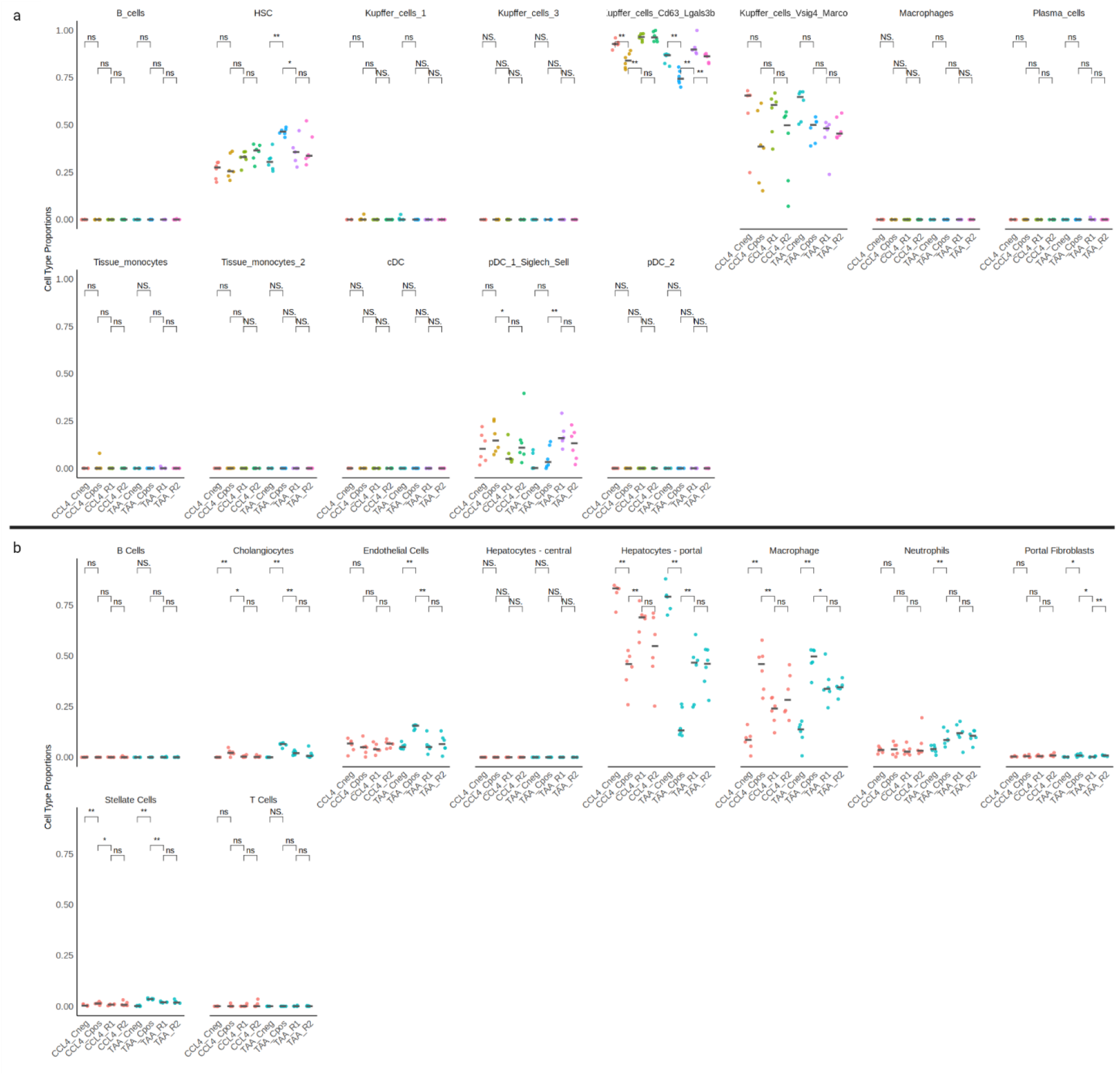
Comparison of all predicted cell populations. **(a)** Cell types obtained from single-cell reference (Ramachandran et al., Nature 2019). In the original dataset, the NPC fraction relates to cells expressing HSC and endothelial markers that persisted through CD45+ cell sorting. **(b)** Cell types obtained from single-nuclei reference (Nault et al., Toxicological Sciences, 2022). Ns = non-significant.

**Supplementary Figure 7_3.**
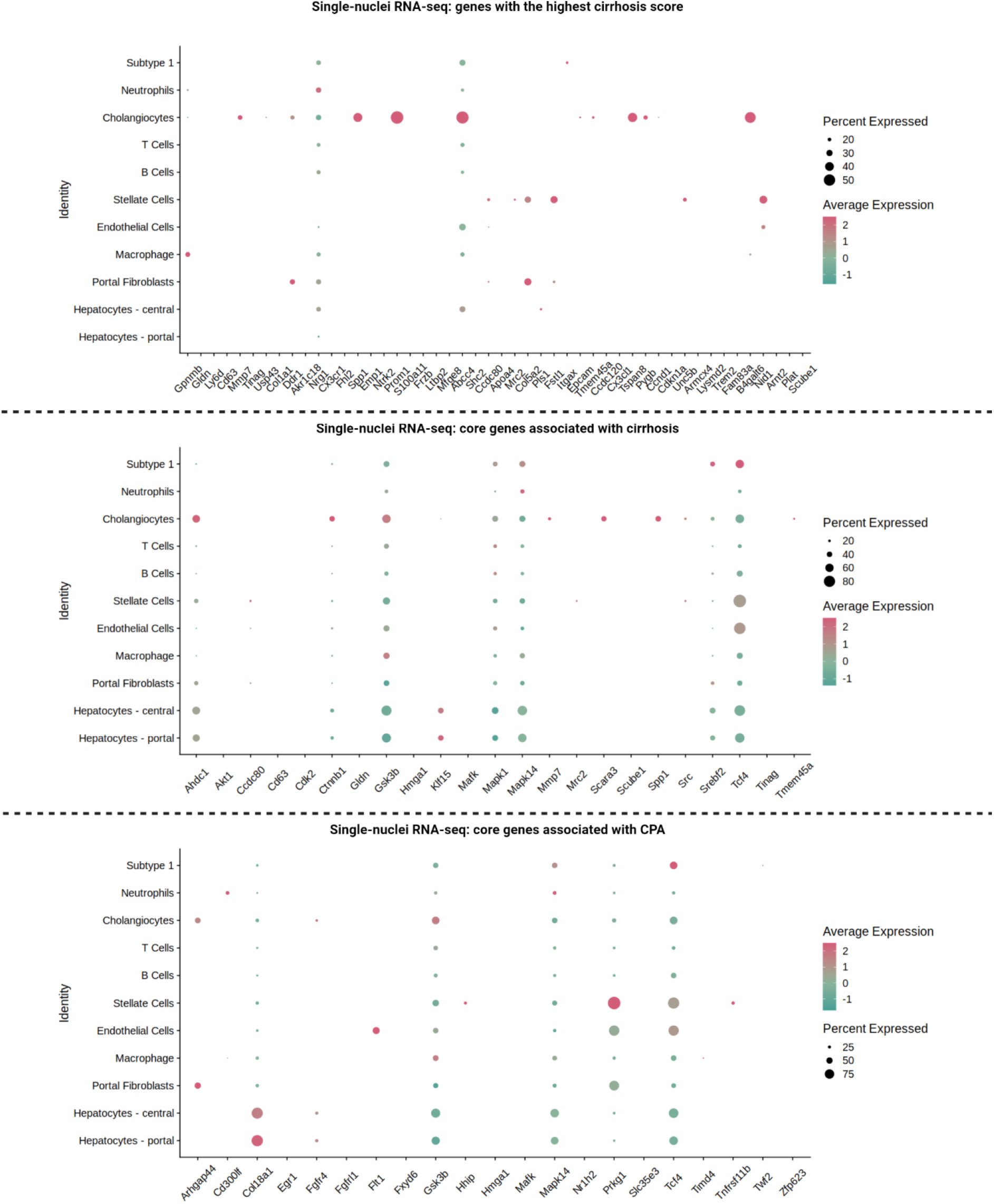
Link of cirrhosis-associated genes to cell types in the single-nuclei RNA-seq dataset. Each row represents a gene set: genes with the highest perturbation (cirrhosis) score; core genes associated with cirrhosis; core genes related to collagen proportionate area, accordingly. Colors indicate expression change from downregulation (green) to upregulation (red). The dot size illustrates the proportion of cells for each specific cell type expressing a gene. Genes from these lists not expressed in either of the cell types are not present. CPA = collagen proportionate area.

**Supplementary Figure 7_4.**
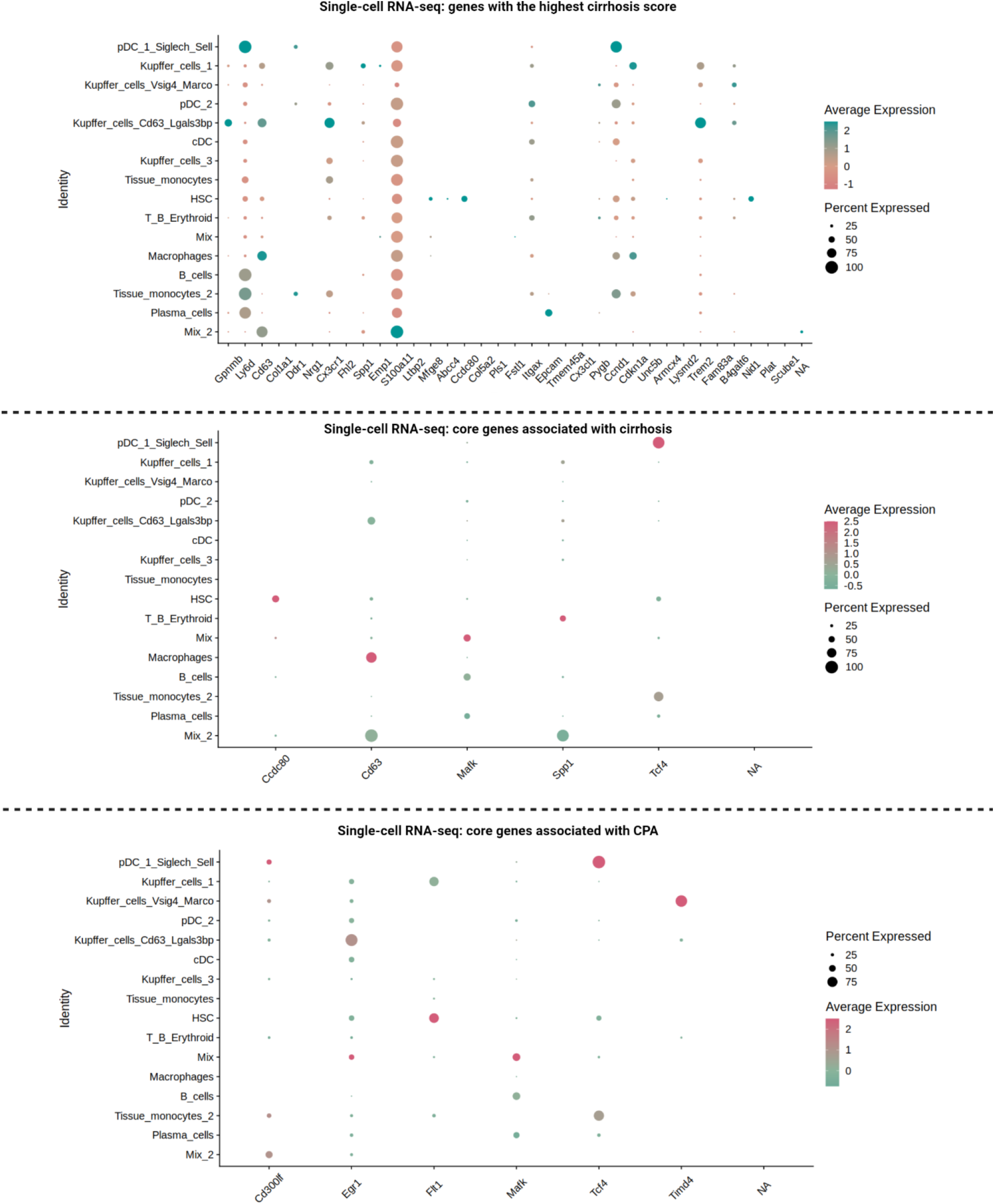
Link of cirrhosis-associated genes to cell types in the single-cell RNA-seq dataset. Each row represents a gene set: genes with the highest perturbation (cirrhosis) score; core genes associated with cirrhosis; core genes related to collagen proportionate area, accordingly. Colors indicate expression change from downregulation (green) to upregulation (red). The dot size illustrates the proportion of cells for each specific cell type expressing a gene. Genes from these lists not expressed in either of the cell types are not present. CPA = collagen proportionate area.

**Supplementary Table 1.** Classification using prioritized biomarkers on human RNA-seq datasets.

| Gene set | Gene set size | Random forest model accuracy | Kappa value | Classification task | Dataset |
| --- | --- | --- | --- | --- | --- |
| *CPZ, ARHGAP44, SRC, PRKG1, ISLR, SPP1, TCF4* | 7 | 0.710 | 0.348 | F0 vs F3/F4 | PRJEB27201 |
| *CPZ, HMGA1, IRF7, MRC2* | 4 | 0.933 | 0.867 | Cirrhotic PH vs healthy control | GSE171248 |
| *CPZ, IGFBP7, SPP1, RCN3, CCDC80, MMP7, KLF15, MRC2* | 8 | 0.761 | 0.495 | F0 vs F3/F4 | GSE225740 |

PH = portal hypertension.

## Notes

https://github.com/xander-p/CDL_regression

### Competing Interest Statement

The group of TR (OP, PK, KBr, BSH, KBa, BS, PS) received funding from the Austrian Federal Ministry for Digital and Economic Affairs, the National Foundation for Research, Technology and Development, Boehringer Ingelheim, and the Christian Doppler Research Association. AFR is supported by Angelini Ventures S.p.A. Rome, Italy.

